# P2Y14 receptor agonist impairs, and antagonist improves whole-body glucose homeostasis and liver steatosis

**DOI:** 10.64898/2026.09.01.748648

**Authors:** Asmita Pramanik, Siva Hariprasad Kurma, Zhiwei Wen, Hongyi Cai, Pierre-Christian Violet, Peter J. Walter, Tamar Demby, Oksana Gavrilova, Srinivas Pittala, Haressh Sajiir, Regina Umarova, Allison Wing, Yaron Rotman, Jürgen Wess, Kenneth A. Jacobson

## Abstract

Knockout of the UDP-glucose-activated P2Y_14_ receptor (P2Y_14_R) in adipocytes or whole-body has been reported to provide metabolic benefits in obese mice. We hypothesized that selective P2Y_14_R activation would lead to metabolic impairments, whereas pharmacological antagonism would improve metabolic deficits in diet-induced obese (DIO) mice. Here, we investigated the metabolic effects of a synthetic P2Y_14_R agonist, UDP-like methylene-bridged MRS2905. Acute P2Y_14_R activation with MRS2905 triggered a robust and prolonged hyperglycemic effect in lean and obese mice with impaired glucose homeostasis. Moreover, MRS2905 treatment of obese mice lowered fasting plasma insulin and increased glucagon levels, along with upregulation of liver JNK phosphorylation and the expression of rate-limiting gluconeogenic genes. The MRS2905-induced hyperglycemic effect was blunted in whole-body P2Y_14_R knockout mice compared with wild-type control. In contrast, a potent P2Y_14_R antagonist mono-ester prodrug (MRS4779) partially reversed the agonist-induced hyperglycemia and restored proper glucose homeostasis after acute treatment. Chronic MRS4779 administration in DIO mice reduced fat mass and improved various metabolic parameters including liver steatosis. Additionally, we examined the roles of P2Y_14_R in hepatocytes of DIO mice. Here, we report that P2Y_14_R was upregulated in liver and hepatocytes from obese mice compared to lean mice. Overnight fasting also upregulated hepatic P2Y_14_R expression. P2Y_14_R deletion from hepatocytes in DIO mice improved fasting blood glucose level and lipid metabolism without improving glucose homeostasis. These results suggest a novel P2Y_14_R function in hepatic lipid metabolism, and P2Y_14_R antagonists may prove useful for the treatment of obesity and obesity-related metabolic disorders.

## 1. Introduction

G protein-coupled receptors (GPCRs) expressed in various metabolic tissues are involved in the regulation of whole-body glucose and energy homeostasis. Dysregulation of glucose homeostasis contributes to hyperglycemia in obesity and type 2 diabetes (T2D) [1, 2]. Thus, GPCR modulators have emerged as important therapeutic targets for anti-obesity and anti-diabetic drug candidates [3–6]. Current treatment regimens, while providing extraordinary benefits, are often associated with adverse side effects, including gastrointestinal disturbances and muscle and bone mass loss, underscoring the need for novel therapeutic targets [7–9]. Additionally, identifying novel drug candidates to improve the metabolic deficits caused by obesity is of great clinical relevance as a standalone candidate or combination therapy.

The P2Y_14_ receptor (P2Y_14_R, formerly GPR105) belongs to the purinergic GPCR family and is activated by nucleotide-sugars, most potently with uridine-5′-diphosphoglucose (UDP-G) [10]. P2Y_14_R couples to G_i_-type G proteins and is widely expressed, including in placenta, adipose tissue, stomach, intestine, and discrete brain regions [11]. This receptor is also expressed in multiple metabolically relevant tissues in both human and mouse, including adipose tissue [6] and pancreatic islets [12]. Interestingly, P2Y_14_R tissue expression is altered in obese individuals with insulin resistance (IR), suggesting that P2Y_14_R may represent a potential therapeutic target for the treatment of obesity and T2D [12]. Ablation of this receptor from macrophages and whole body improves insulin sensitivity in diet-induced obese (DIO) mice [13]. Moreover, a recent study showed P2Y_14_R expression in human and mouse adipocytes, and that overnutrition causes P2Y_14_R upregulation in adipose tissue [14]. P2Y_14_R activation inhibits lipolysis in white adipose tissues, and adipocyte-specific P2Y_14_R knockout mice show reduced body weight and improved whole body glucose homeostasis under DIO conditions by increasing fasting lipolysis [14]. P2Y_14_R activation with endogenous ligand (UDP-G) in mouse islets decreases glucose-stimulated insulin secretion (GSIS) [12]. Activation of P2Y_14_R on smooth muscle promotes gastrointestinal emptying, and P2Y_14_R deficiency in pancreatic β-cells reduces glucose tolerance and insulin release [15]. P2Y_14_R expression in liver and hepatic stellate cells (HSCs) was upregulated in a CCl_4_-induced liver fibrosis mouse model, which was associated with UDP-G release from dying hepatocytes causing activation of HSCs. P2Y_14_R deletion from HSCs selectively attenuates liver fibrosis in mouse [16]. Despite this finding, its role in hepatocytes in the context of overnutrition or fasting-induced stress has not been explored.

At high concentrations (µM range), UDP-G can also activate the P2Y_2_R and P2Y_6_R [17]. To study P2Y_14_R under physiological and pathophysiological conditions, we therefore used highly selective, synthetic P2Y_14_R ligands. Specifically, we employed a UDP analogue, MRS2905, and UDP-G analogue, MRS2690, two potent and selective synthetic P2Y_14_R agonists [17], to investigate the metabolic effects triggered by P2Y_14_R activation in both lean and obese mice. We also tested the metabolic effects of P2Y_14_R blockade by using an antagonist mono-ester prodrug, MRS4779 [18] in DIO mouse models. Moreover, we generated the first hepatocyte-specific P2Y_14_R knockout mouse model to study the hepatic functions of P2Y_14_R in obese mice.

Our data show that the acute P2Y_14_R activation in lean and obese mice impaired whole body glucose homeostasis by decreasing plasma insulin and increasing glucagon levels. In contrast, acute blockade of P2Y_14_R signaling partially blunted MRS2905-induced hyperglycemia. Chronic MRS4779 treatment in DIO mice reduced fat mass, without alteration of lean mass, and showed trends to improve metabolic parameters in obese mice. We establish that nutrient-induced stress alters P2Y_14_R expression in liver and hepatocytes, but hepatocyte-selective P2Y_14_R deficiency did not improve glucose homeostasis in DIO mice. However, deletion of P2Y_14_R in hepatocytes improved steatosis and lipid metabolism in obese mice. These findings suggest that blocking P2Y_14_R signaling may prove a useful novel approach for the treatment of obesity and obesity-related metabolic disorders.

## 2. Materials and methods

### 2.1. Materials

P2Y_14_R agonists UDP-glucose disodium salt (Abcam), MRS2690 disodium salt (Bio-Techne), and MRS2905 trisodium salt (Bio-Techne) were purchased commercially. The P2Y_14_R highly selective antagonist PPTN-HCl [4-[4-(4-piperidinyl)phenyl]-7-[4-(trifluoromethyl)phenyl]-2-naphthalenecarboxylic acid hydrochloride] was purchased from Bio-Techne. The other two P2Y_14_R antagonists, MRS4738 and MRS4779 (trifluoroacetate salt), were synthesized as previously reported [18]. UDP-glucose, MRS2690, and MRS2905 were dissolved in sterile 0.9% saline for in vitro or in vivo experiments. Stock solutions of PPTN-HCl and MRS4738 were prepared in DMSO and diluted in 1X PBS for desired concentration. For in vivo experiments, MRS4779 was first dissolved in DMSO (Sigma-Aldrich), followed by the sequential addition of Kolliphor EL (Sigma-Aldrich) and sterile saline (DMSO/Kolliphor EL/Saline, 5-10: 5-10: 80-90% (v/v)).

### 2.2. Mouse maintenance and diet

Wild-type (WT) C57BL/6NTac (C57BL/6N, Taconic Biosciences), adipocyte-*P2ry14^-/-^ (adipo-P2ry14^-/-^)*, whole body *P2ry14 ^-/-^(WB-P2ry14 ^-/-^)*, and hepatocyte-*P2ry14 ^-/-^ (Hep-P2ry14 ^-/-^)*, mice were maintained on feed and water provided ad libitum with a 12 h light/ dark cycle (lights off at 6 PM and on at 6 AM) at room temperature (23°C), and all experiments were conducted in group-housed mice unless indicated otherwise. Unless indicated otherwise, mice were maintained on a regular chow (RC) diet (LabDiet, cat. no. 5018; energy density, 3.05 kcal/g). A subset of male mice was fed a high-fat diet (HFD) (60% kcal fat; F3282, Bio-Serv, energy density 5.49 kcal/g) for at least 8 weeks starting at 6 weeks of age to generate diet-induced obese (DIO) mice, and body weight was measured weekly. Metabolic tissues were collected from mice in the experimental groups after euthanasia by CO₂ asphyxiation. All animal procedures were approved by the Institutional Animal Care and Use Committee (IACUC, K083-LBC-20 dated 09.01.2020 and K083-LBC-23 dated 09.01.2023) of the National Institute of Diabetes and Digestive and Kidney Diseases (NIDDK) and conducted in accordance with the U.S. National Institutes of Health (NIH) Guidelines for Animal Research. The patient samples were part of an approved NIH clinical protocol (#19DK0072), and all participants or their legal representatives granted written informed consent to take part in the study and consent to publish.

### 2.3. Mouse models

Mice lacking P2Y_14_R selectively in adipocytes were generated according to a previous study [19]. Briefly, *P2ry14^fl/fl^* mice (NIEHS, NIH, Research Triangle Park, North Carolina, genetic background: C57BL/6J) were crossed with adipoq-Cre mice expressing recombinase under the control of the adiponectin promoter (Jackson Laboratories; stock no. 010803; genetic background: C57BL/6J). Mice used for experiments, *P2ry14^fl/fl^*(control), and adipoq-*Cre P2ry14^fl/fl^* (adipo-*P2ry14 ^-/-^*), were littermates. Global P2Y_14_R knockout mice were generated by crossing *P2ry14^fl/fl^* mice with CMV-Cre mice expressing recombinase under the transcriptional control of a human cytomegalovirus minimal promoter (Jackson Laboratories; stock# 006054, genetic background: C57BL/6J) [14]. Throughout the text, we refer to these mice as WB-*P2ry14^-/-^* mice. We also generated mice in which we disrupted the *P2ry14* gene specifically in hepatocytes (Hep-*P2ry14*^-/-^). These mice were obtained by crossing *P2ry14^fl/fl^* mice with albumin-Cre mice (Jackson Laboratories; stock no. 003574; genetic background: C57BL/6J).

### 2.4. In vivo pharmacokinetics study

To measure the time-dependent plasma concentrations of P2Y_14_R agonists, 12-week-old male C57BL/6NTac mice were injected intraperitoneally (i.p.) with UDP-G or MRS2905 (10 mg/kg each). At defined post-injection time points, blood (∼25 μl) was collected from the tail vein into heparinized capillary tubes, centrifuged at 2000Xg for 10 min at 4°C, and the plasma was analyzed by UPLC-MS/MS. A detailed UPLC-MS/MS method for detecting and quantifying UDP-glucose and MRS2905 is described in Supplementary Information. Plasma concentrations of each agonist in a time-dependent manner were analyzed to derive C_max_ (peak y) and T_max_ (peak x) values.

### 2.5. Body composition analysis

Body composition (lean vs. fat mass) was determined in non-anesthetized mice using EchoMRI100H analyzer (EchoMRI LLC).

### 2.6. Metabolic studies

Various metabolic tests were performed with male WT, adipo-*P2ry14^-/-^*, WB-*P2ry14 ^-/-^* and both male and female Hep-*P2ry14^-/-^* mice (age: 10-24 weeks). Intraperitoneal glucose tolerance tests (ipGTT) were conducted in mice fasted for 12-14 h. Mice consuming RC diet received a glucose dose of 2 g/kg body weight, while HFD-fed mice were injected with 1 g/kg glucose, respectively. Blood glucose levels were measured prior to glucose injection and at defined post-injection time points. Blood glucose measurements were carried out with a glucometer (Contour Next EZ, Bayer) using blood obtained from the tail vein. Insulin tolerance tests (ITT) and glucagon challenge tests (GCT) were performed after a 5 h fast. For ITT, chow-fed lean mice or HFD-fed obese mice were injected i.p. with 0.75 U/kg or 1.0 U/kg of human insulin (HumulinR, Eli Lilly), respectively. For GCT, lean or obese mice received a glucagon dose of 16 µg/kg or 20 µg/kg (i.p., Sigma-Aldrich), respectively. For pyruvate tolerance tests (PTT), mice were fasted overnight (12-14 h), and then sodium pyruvate (2.0 g/kg, or 1.5 g/kg, i.p., Sigma-Aldrich) was injected, followed by monitoring of blood glucose levels at indicated time points. Area of the curve (AOC) values for all metabolic tests were obtained by subtracting the baseline values in Prism (v.10.2.2, GraphPad) as recommended when baseline values differ between experimental groups [20].

### 2.7. Plasma metabolic profiling

Blood was collected from the mouse tail vein in chilled K_2_-EDTA containing tubes (Sarstedt). Blood was centrifuged at 4°C for 10 min at 10,000Xg to obtain plasma. ELISA kits (Crystal Chem) were used to measure plasma insulin and glucagon levels, following the manufacturer’s instructions. Plasma free fatty acid (FFA) levels were determined using a commercially available kit (Fujifilm, Wako Diagnostics). Plasma triglyceride and cholesterol were measured using commercially available kits (T7532-120, Pointe Reagents; C5710-120, Fisher Scientific).

### 2.8 Quantification of glycogen and triglyceride in liver

The liver tissue samples (∼50 mg) were homogenized in ice-cold 25 mM citrate buffer, pH 4.2, containing 2.5 g/L NaF. To remove debris, homogenates were centrifuged at 12000 rpm for 5 min at 4°C, and supernatant was used for quantification of glycogen using a Glycogen assay kit (Abnova). The glycogen content was normalized by liver tissue weight.

Total lipid was extracted by homogenizing frozen liver tissue (∼50 mg) in phosphate-buffered saline followed by Folch extraction method. A mixture of chloroform/methanol (2:1) was added to the liver homogenate and vortexed vigorously to extract lipid. The homogenate was centrifuged at 2500 rpm for 10 min at 4°C, and the lipid-containing organic phase was collected and dried overnight by Speed Vac (Savant). Total lipid content was dissolved in ethanol containing 1% Triton X-100 (Sigma-Aldrich). Triglyceride levels were quantified spectrophotometrically using a L-Type Triglyceride M kit (Fujifilm Wako Diagnostics) according to the manufacturer’s instructions and normalized to liver weight.

### 2.9. Acute activation with P2Y_14_R agonists

To investigate the metabolic effects of acute P2Y_14_R activation, its endogenous agonist (UDP-G) or highly selective P2Y_14_R agonists (MRS2690 and MRS2905) were injected i.p. (10 mg/kg, equivalent to 16.4, 16.0 and 20.7 µmol/kg, respectively) in overnight fasted or fed WT (lean and obese) mice. Fasted or fed blood glucose levels were measured at different time points after administration. For effective dose determination, MRS2905 was administered in obese overnight-fasted mice with three doses (10 mg/kg, 5 mg/kg, and 1 mg/kg, i.p.), and the fasting blood glucose level was measured at different time points. MRS2905 (10 mg/kg, i.p.) was also administered to HFD control, adipo-*P2ry14*^-/-^, and lean WB-*P2ry14*^-/-^ mice, followed by the determination of fasting blood glucose levels.

### 2.10. Acute effect of antagonist prodrug MRS4779

To examine the acute effects of blocking P2Y_14_R signaling, obese WT mice were treated with P2Y_14_R antagonist prodrug MRS4779 (10 mg/kg, i.p., equivalent to 14.6 µmol/kg), either alone or together with MRS2905 or vehicle (DMSO/Kolliphor EL/Saline, 10:10:80 (v/v) for MRS4779 and 0.9% sterile saline for MRS2905, in the evening 1-2 h before food removal. On the following morning, mice were injected with a second MRS4779 dose (10 mg/kg, i.p.). After 2 h, mice were treated with the P2Y_14_R agonist MRS2905 (10 mg/kg, i.p.) or saline. Fasting blood glucose was measured in tail vein blood immediately before (0 min) and 45 min after agonist injection. GTT, ITT, and GSIS were also performed in vehicle- and MRS4779-only treated groups.

### 2.11. Chronic treatment with antagonist prodrug MRS4779

In vivo efficacy of prodrug antagonist MRS4779 was studied in DIO mice. Male WT mice were fed with a HFD for 12 weeks to reach 45 g of body weight. Mice were single-housed and baseline parameters (body weight, fed blood glucose levels and food intake) were recorded for 3 days with daily intraperitoneal injections of vehicle. Starting from day 0 designated groups received either vehicle (DMSO/Kolliphor EL/Saline, 5:5:90 (v/v)) or MRS4779 (5 mg/kg, i.p., dissolved in vehicle) twice daily for 10 days. In a similar fashion, age-matched lean mice (RC-control group) received two doses of vehicle daily. Fed blood glucose, food intake, and body weight were monitored in the morning daily. On days 11 and 13, ITT and GTT were performed, respectively. On day 14, mice were euthanized via CO_2_. Different tissues were removed and weighed, and samples were either snap-frozen or fixed in buffered 10% neutral formalin. Blood was collected by cardiac puncture to determine the levels of hormones and metabolites.

### 2.12. RNA extraction and gene expression analysis

Liver tissue from different experimental mice was collected and snap frozen in liquid nitrogen. Total liver RNA was extracted using the RNeasy Mini Kit (Qiagen). Total RNA was extracted from treated and untreated AML12 mouse hepatocyte cells with phenol-chloroform method (TriPure isolation solution, Roche). RNA concentration was measured by Nanodrop (Thermo Scientific Nanodrop 2000), and 500 ng of RNA was converted into cDNA using High-capacity RNA to cDNA kit (Applied Biosystems). Quantitative PCR was performed using the Fast SYBR green PCR Master Mix (Applied Biosystems). Gene expression was normalized to the expression of *36b4* using the ΔΔCt method. The primer sequences used in this study are provided in Supplementary Table 2.

### 2.13. Western blotting

Liver tissues were homogenized in liver lysis buffer (4% SDS,100 mM Tris pH 8.0, 5mM DTT mM, 0.5 mM EDTA, and 1 mM phenylmethylsulfonyl fluoride) supplemented with EDTA-free protease inhibitor cocktail and phosphatase inhibitors cocktail (Roche). Protein concentrations in the lysates were determined using a BCA Protein Assay Kit (Pierce, Thermo Fisher Scientific). Protein (10 µg) was denatured in NuPAGE LDS sample buffer (Thermo Fisher Scientific) and 5% β-mercaptoethanol at 95°C for 5 min. Protein lysates were separated using 4–12% SDS-PAGE (Invitrogen) and transferred to nitrocellulose membranes (Trans-Blot turbo transfer system, Bio-Rad). The membrane was blocked with PBS blocking buffer (Odyssey) at room temperature (RT) for 1 ah followed by primary antibody incubation in PBS blocking buffer with 0.1% Tween-20 (PBSBBT) overnight at 4°C. The following day, the blot was washed four times (5 min each) with 1X PBS with 0.1% Tween-20 (PBST). The membrane was incubated with a suitable IR-labeled secondary antibody (1:10,000, Odyssey) diluted in PBSBBT at RT for 1 h. The blot was washed four times with 1X PBST and developed under Odyssey CLx infrared imaging system. Band intensity was quantified using Image Studio Lite software (Li-Cor). A list of primary and secondary antibodies used is provided in Supplementary Table 3.

### 2.14. Tissue histology

Adipose tissue and liver were dissected from euthanized mice and fixed in 10% neutral buffered formalin solution. All tissue samples were sectioned and stained using standard staining techniques. H&E- and Oil Red O (ORO)-stained sections were visualized and images captured using a Keyence Microscope (BZ-9000). Adipose tissue sections were analyzed for adipocyte size and number using Image J software with Marcos (http://imagej.nih.gov/ij).

### 2.15. Hepatocytes isolation from mice

Primary hepatocytes were isolated from the livers of RC or HF diet fed male mice using a collagenase perfusion protocol [21]. Approximately 5 × 10^5^ hepatocytes per well were seeded onto collagen I-coated 6-well plates (Corning) and cultured at 37°C in a humidified incubator containing 5% CO₂. Cells were maintained in DMEM (Gibco) with 4.5 g/L glucose and 10% FBS (Gibco). Once the hepatocytes reached approximately 60-70% confluence, they were used for glucose production or gene expression study.

Primary mouse hepatocytes (5 × 10^5^ cells/well) from obese mice were seeded in 6-well collagen-coated plates (Corning) for 4-6 h at 37°C in phenol red-free DMEM supplemented with 10% FBS and 4.5 g/L glucose. The culture medium was subsequently replaced with phenol red-free DMEM containing 1 g/L glucose, and the cells were cultured overnight. The following day, hepatocytes were washed thoroughly with PBS and incubated in glucose- and phenol red-free DMEM supplemented with the gluconeogenic substrates sodium lactate (20 mM) and sodium pyruvate (2 mM). Cells were then incubated for 5 h at 37 °C in the presence of glucagon (100 nM), MRS2905 (10 nM and 100 nM). Following incubation, the culture medium was collected, and glucose concentrations were quantified using a glucose assay kit (Sigma-Aldrich). For normalization to total cellular protein, hepatocytes were lysed directly in the wells using RIPA buffer supplemented with a protease inhibitor cocktail (Roche), and protein concentrations were determined using the BCA assay.

In another experiment, primary hepatocytes were isolated from lean and obese WT mice. Gene expression study was performed to compared expression of *P2ry14* by RT-qPCR in hepatocytes isolated from liver of lean and obese mice. The primer sequences are provided in Supplementary Table 2.

### 2.16 Cell culture and palmitic acid treatment

The AML12 cell line was obtained from the American Type Culture Collection (ATCC CL-2254). AML12 cells were grown in a DMEM/F-12 (1:1) media (Gibco) and supplemented with 10% FBS (Gibco), 1% penicillin-streptomycin (Gibco), 1% Insulin-transferrin-sodium selenite media supplement (Gibco) and 100 nM dexamethasone (Sigma-Aldrich) and maintained at 37°C with a 5% CO_2_ atmosphere. AML12 cells were exposed to 500 μM palmitic acid (PA) (Sigma-Aldrich) conjugated to fatty acid-free bovine serum albumin (BSA) (Sigma-Aldrich) for 24, 48, and 72 h. Cells treated with BSA were used as controls. To study effect of P2Y_14_R antagonists (PPTN and MRS4738) in PA induced steatosis model, cells were co-incubated with PA (500 µM) and PPTN (100, 200, and 400 nM) or MRS4738 (100, 250 and 500 nM) for 48 h. Cells were stained with ORO stain (Sigma-Aldrich) to detect lipid droplets. Images were captured with a Keyence 7000 microscope (20X).

### 2.17 MTT assay

Cell viability was assessed in AML12 cultures treated with PPTN (0.1-10 µM) and MRS4738 (0.1-10 µM) for 48 h. 0.1% DMSO was used as a solvent control. Cells were incubated with MTT (3-(4,5-dimethylthiazol-2-yl)-2,5-diphenyl-2H-tetrazolium bromide, Sigma-Aldrich) for 4 h, and formazan crystal was dissolved with dissolving solution (Sigma-Aldrich). Absorbance was measured at 560 and 630 nm using a microplate reader (BMG Labtech), and the percentage of cell viability was calculated, use following formula.

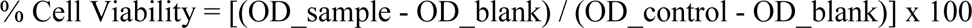

### 2.18 Human subjects and sample collection

Data were analyzed from human subjects with metabolic dysfunction-associated steatotic liver disease (MASLD, previously termed NAFLD) that were included in a clinical trial (clinicaltrials.gov NCT03884075). Briefly, adult subjects were included if they had histologically confirmed MASLD with liver fat content ≥10% by ^1^H-MR spectroscopy, and average alcohol consumption < 30 g/d (men) or < 20 g/d (women). Major exclusion criteria included decompensated liver disease, major comorbidities and uncontrolled diabetes. All subjects met current definition of MASLD [22]. The study was approved (#19DK0072) by the National Institutes of Health Institutional Review Board, and all participants provided written informed consent.

After an overnight fast, a subcutaneous adipose tissue sample was obtained under local anesthesia using a 10G Spirotri suction cannula. This was followed immediately by a percutaneous liver biopsy under local anesthesia and conscious sedation using a 16G cutting needle. Liver and adipose samples were aliquoted immediately after procurement, flash-frozen at the bedside, and stored in -80°C until analysis. Liver histology was scored by an expert hepatopathologist, according to the NASH-CRN score [23].

A 2-hour oral glucose tolerance test was performed two days prior to liver and adipose biopsies. After an overnight fast, subjects ingested 75 g of glucose in 300 ml of water, with blood samples for glucose and insulin collected at baseline and every 30 minutes. Insulin sensitivity and beta-cell function were estimated according to the homeostasis model assessment (HOMA) from fasting values, using HOMA-IR and HOMA-B, respectively.

### 2.19 Human RNA sequencing and analysis

Total RNA was extracted from human adipose tissue using TRIzol (Thermo Fisher Scientific), followed by purification with the Qiagen RNeasy Mini Kit according to the manufacturer’s instructions, including on-column DNase I digestion. Tissue was bead-homogenized (Lysing Matrix D) in TRIzol, and the lipid layer was removed prior to column purification. RNA was quantified by NanoDrop and stored at -80°C.

RNA quality was assessed using a NanoDrop spectrophotometer, and an Agilent 2100 Bioanalyzer. Libraries were sequenced on an Illumina NovaSeq 6000 system to generate paired-end 150 bp reads. Raw sequencing data were quality-assessed using FastQC (Babraham Institute, http://www.bioinformatics.babraham.ac.uk/projects/fastqc/), and reads were processed by removing low-quality bases and adapter sequences using Trimmomatic (v0.27) [24]. Quality-filtered reads were aligned to the Ensembl human reference genome (release 114) using the STAR aligner (v2.7.11b) with two-pass mapping [25]. Uniquely mapped reads were quantified using RSEM (v1.3.3) [26], and gene counts were normalized using the median-of-ratios method implemented in the DESeq2 R package (v1.48.2) [27]. Expression of P2ry14 was correlated to clinical measures using Pearson correlations with gender as a covariate.

### 2.20. Statistical analysis

Statistical analyses were performed using GraphPad Prism 10.2.2 (GraphPad Software). Data are expressed as mean ± SEM for the number of observations indicated in the figure legends. Data were tested by One-way or Two-way ANOVA, followed by Sidak’s multiple comparison test tests, or by 2-tailed unpaired Student’s t test as appropriate. A P value <0.05 was considered statistically significant.

## 3. Results

### 3.1. Acute P2Y_14_R activation causes hyperglycemia and impairs whole-body glucose homeostasis in lean and obese WT mice

To study the effect of acute P2Y_14_R activation on blood glucose levels, we administered UDP-G, the endogenous P2Y_14_R agonist, and two P2Y_14_R-selective, synthetic agonists (MRS2690 and MRS2905) to lean and obese WT male mice. Interestingly, treatment of lean mice with a single dose of UDP-G (10 mg/kg, i.p.) did not lead to any changes in fasting blood glucose levels, as compared to vehicle-injected control mice (Supplementary Fig. S1A, B). However, i.p. treatment with either MRS2690 or MRS2905 (10 mg/kg, i.p.) significantly increased fasting blood glucose levels under the same experimental conditions but to different extents (Supplementary Figs. S1, Fig. 1B). MRS2905 was selected for further studies as it showed a robust effect in this vivo model. We next measured plasma levels of insulin and glucagon, which play key roles in maintaining euglycemia. Treatment of lean mice with MRS2905 (10 mg/kg, i.p.) had no significant effect on plasma insulin levels but resulted in significantly increased plasma glucagon levels at 30 min (Figs. 1C, D). MRS2905 treatment did not induce any changes in blood glucose or plasma insulin levels in freely fed lean mice (Figs. 1E, F)

**Figure 1:**
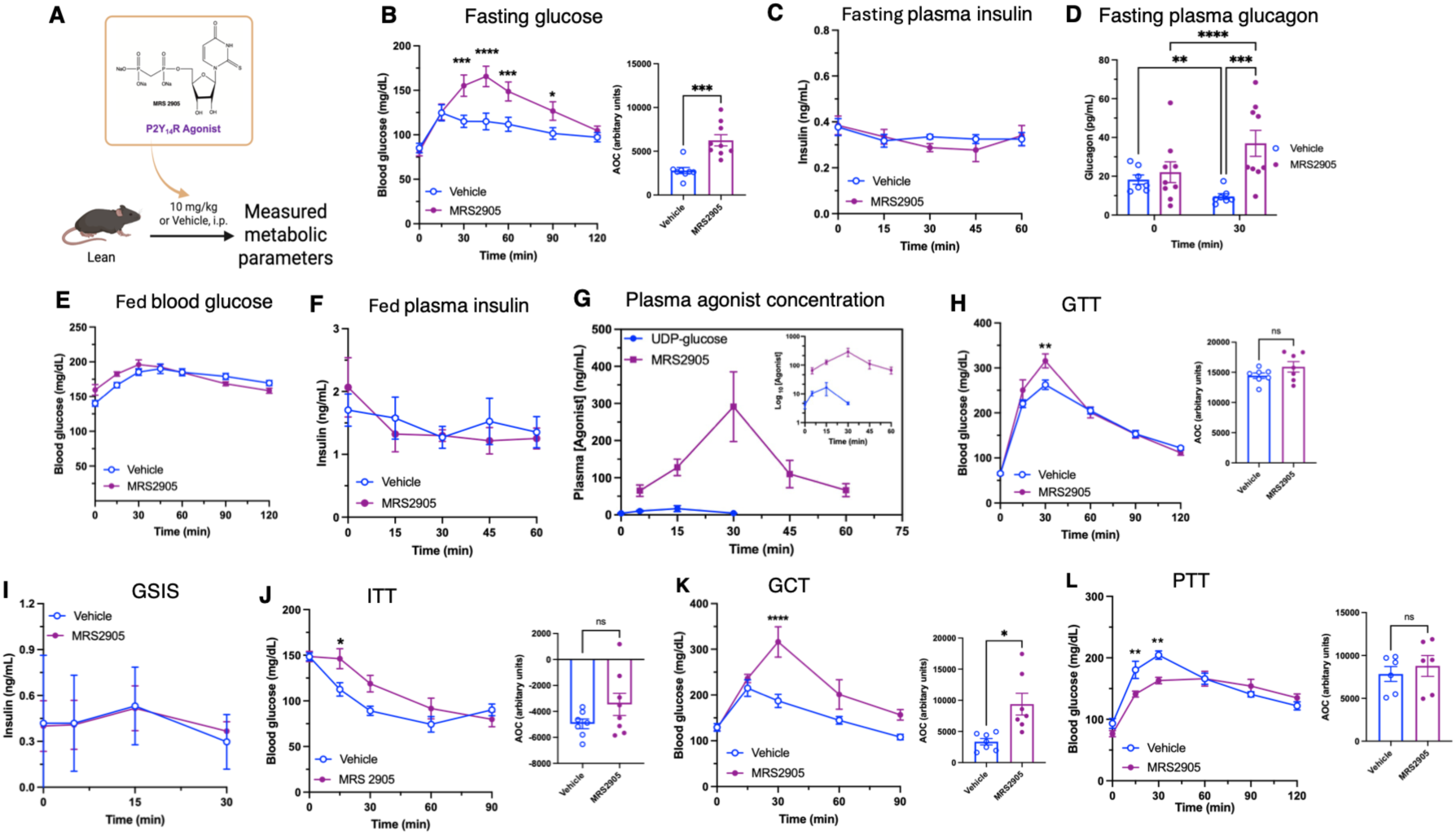
Acute activation of P2Y_14_R increases fasting blood glucose and alters pancreatic endocrine hormones with impairing glucose metabolism in lean mice maintained on regular chow (RC) diet. **(A)** Schematic presentation of metabolic parameter studies in lean mice. **(B)** Overnight fasting blood glucose levels in lean mice injected saline or MRS2905 (n=8-9/group, 10 mg/kg, i.p.) at 0, 15, 30, 45, 60, 90 and 120 min. **(C)** Overnight fasting plasma insulin levels in saline or MRS2905 (n=4/group) treated mice at indicated time points. **(D)** Overnight fasting plasma glucagon levels in saline or MRS2905 (n=7-9/group) treated mice at before and 30 min after MRS2905 administration. **(E)** Blood glucose and **(F)** plasma insulin levels in lean mice fed freely and treated with saline or MRS2905 (n=5/group) at indicated timepoints. **(G)** Pharmacokinetic measurements of UDP-glucose and MRS2905 in 12-week-old male C57BL/6NTac mice. UDP-glucose and MRS2905 (n=3-5/group) concentration in plasma as a function of time after single intraperitoneal dose (10 mg/kg) in C57BL/6 NTac male mice. The inset shows the data plotted with the y-axis in log_10_ scale and x-axis =time (0-60 min). **(H)** I.p. glucose tolerance test performed after co-administration of saline or MRS2905 (10 mg/kg, ip) with glucose bolus (n=7/group, 2g/kg, i.p.), **(I)** Glucose-stimulated insulin secretion measured after co-administration of saline or MRS2905 (10 mg/kg, ip) with glucose bolus (n=6-8/group, 2 g/kg, i.p.), **(J)** Insulin tolerance test carried out with co-administration of saline or MRS2905 (10 mg/kg, ip) with insulin (n=7-8/group, 1U/kg, i.p.), **(K)** Glucagon challenge test performed with co-administration of saline or MRS2905 (10 mg/kg, ip) with glucagon (n=7/group, 16 µg/kg, i.p.), and **(L)** Pyruvate tolerance tests carried out with co-administration of saline or MRS2905 (10 mg/kg, ip) with pyruvate (n=6/group, 2 g/kg, i.p.). AOC (area of the curve) is presented along with each metabolic test. Data are given as the means ±SEM. *P < 0.05, **P < 0.01, ***P < 0.001, and ****P < 0.0001, as compared with the corresponding saline treated control group [Two-way ANOVA followed by Sidak’s multiple comparison test tests, or by 2-tailed unpaired Student’s t test as appropriate].

We next studied the pharmacokinetic (PK) properties of UDP-G and MRS2905 following i.p. administration (10 mg/kg) to male C57BL/6NTac mice. The plasma concentrations of the two agonists were quantified at different time points after i.p. injection using UPLC-MS/MS analysis. Chromatograms, retention time and *m/z* transition for UDP-G and MRS2905 are shown in supplementary information (Supplementary Fig. S3 and Table S1) and analytes in plasma were quantified. As UDP-G is an endogenous metabolite, its mean plasma concentration at 0 min was 4.1±1.1 ng/mL whereas the maximum plasma UDP-G concentration was reached at 15 min (T_max_ = 15 mins, C_max_= 16.6±8.1 ng/mL) after a single i.p. dose and returned to baseline at 30 min (Fig. 1G and inset figure). In contrast, MRS2905 attained a maximum concentration (C_max_= 291.5±94.0 ng/mL) that was approximal 18 times higher than UDP-G and displayed a T_max_ at 30 min (Fig. 1G and inset figure). Pharmacokinetics data suggested that MRS2905 is metabolically stable compared to endogenous agonist UDP-G in vivo.

In obese mice, a single dose of UDP-G (10 mg/kg, i.p.) or MRS2690 or MRS2905 significantly increased fasting blood glucose levels, as compared to vehicle-treated control mice with different levels. UDP-G and MRS2690 induced hyperglycemia is comparable and in lesser extent compared to MRS2905 induced hyperglycemia (Supplementary Figs. S1C). MRS2905 treatment of obese mice (10 mg/kg, i.p.) led to even stronger hyperglycemic effects (Fig. 2B). A dose-response curve for MRS2905 was performed in obese mice to identify an optimal efficacious dose. We found that 10 mg/kg, i.p. of MRS2905 caused a significant increase of fasting blood glucose (Supplementary Fig. S1D, E). Based on these findings, we selected MRS2905 for more detailed metabolic studies. However, MRS2905 (10 mg/kg) treatment of obese mice led to decreased plasma insulin but increased plasma glucagon levels at 30 min after an overnight fast (Figs. 2C, D). Similarly, MRS2905 caused pronounced hyperglycemia and reduced plasma insulin levels in freely fed obese mice (Figs. 2E, F).

**Figure 2:**
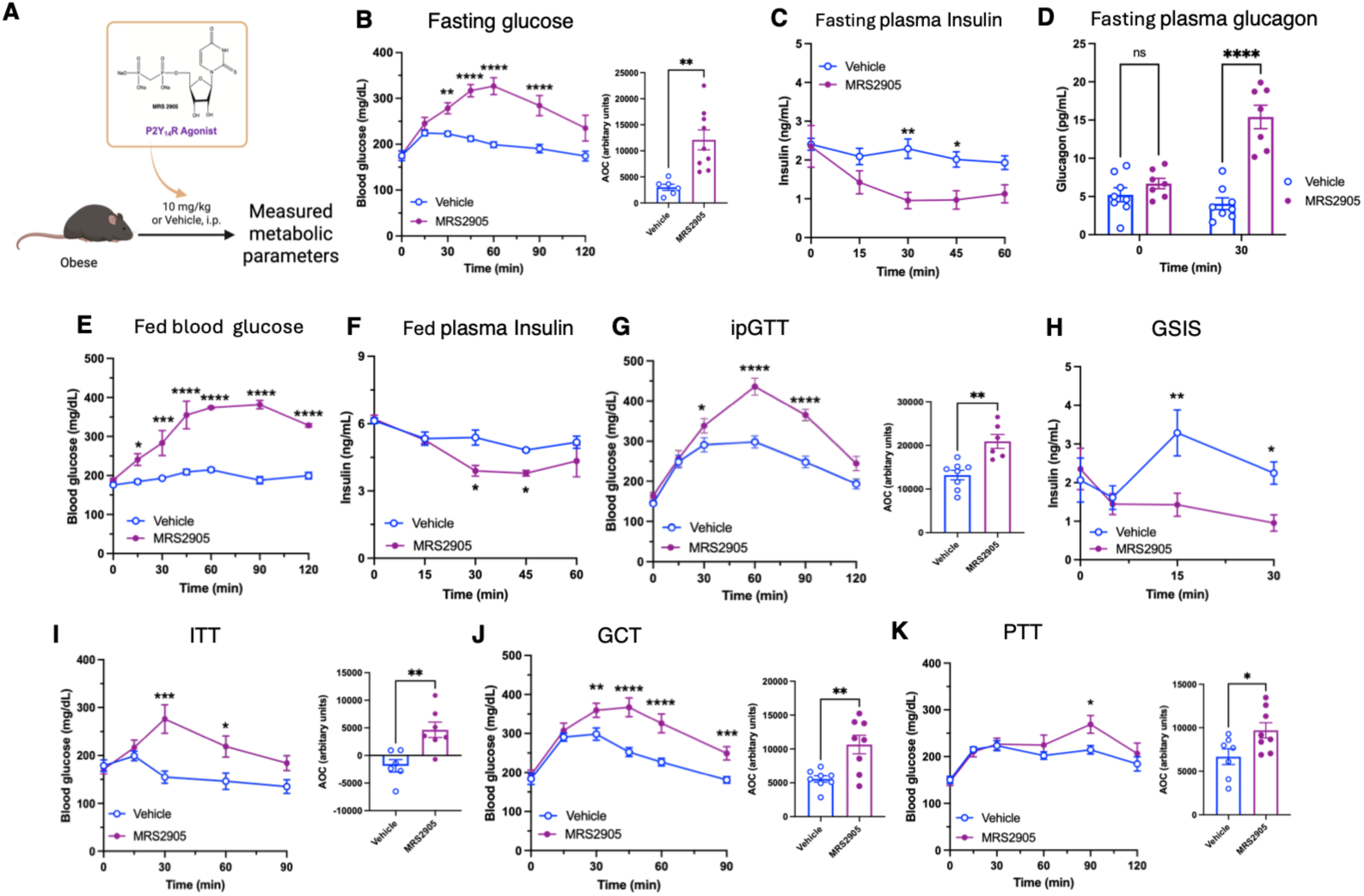
Acute activation of P2Y_14_R in wildtype mice maintained on an obesogenic diet causes hyperglycemia and alters pancreatic endocrine hormones with severe glucose metabolism impairment. **(A)** Schematic presentation of metabolic parameter studies in obese mice. **(B)** Overnight fasting blood glucose levels in obese mice injected saline or MRS2905 (n=6-9/group, 10 mg/kg, i.p.) at 0, 15, 30, 45, 60, 90, and 120 min. **(C)** Overnight fasting plasma insulin levels in saline or MRS2905 (n=6-7) treated obese mice at indicated time points. **(D)** Overnight fasting plasma glucagon levels in saline or MRS2905 (n=3-4/group) treated mice at before and 30 min after MRS2905 administration. **(E)** Blood glucose and **(F)** plasma insulin levels in obese mice fed freely received saline or MRS2905 (n=5-6/group) at indicated timepoints. **(G)** I.p. glucose tolerance test performed after co-administration of saline or MRS2905 (10 mg/kg, ip) with glucose bolus (n=6-8/group, 1.5 g/kg, i.p.) in obese mice, **(H)** Glucose-stimulated insulin secretion measured after co-administration of saline or MRS2905 (10 mg/kg, ip) with glucose bolus (n=6/group, 1.5 g/kg, i.p.), **(I)** Insulin tolerance test carried out with co-administration of saline or MRS2905 (10 mg/kg, ip) with insulin (n=6-7/group, 0.75 U/kg, i.p.), **(J)** Glucagon challenge test performed with co-administration of saline or MRS2905 (10 mg/kg, ip) with glucagon (n=8/group, 20 µg/kg, i.p.), and **(K)** Pyruvate tolerance tests carried out with co-administration of saline or MRS2905 (10 mg/kg, ip) with pyruvate (n=7-8/group, 1.5 g/kg, i.p.). Data are given as the means ±SEM. *P < 0.05, **P < 0.01, ***P < 0.001, and ****P < 0.0001, as compared with the corresponding saline treated control group [Two-way ANOVA followed by Sidak’s multiple comparison test tests, or by 2-tailed unpaired Student’s t test as appropriate].

In lean and obese mice various metabolic tests carried out with treatment of MRS2905 In an i.p. glucose tolerance test (ipGTT), MRS2905 treatment (10 mg/kg i.p.) of lean mice significantly increased blood glucose excursions at the 30 min time point, as compared with saline-injected control mice (Fig. 1H). Somewhat surprisingly, MRS2905 had no significant effect on GSIS in lean mice (Fig. 1I). In contrast, MRS2905 administration led to severely impaired glucose tolerance in obese mice (Fig. 2G). However, in obese mice, MRS2905 caused a decrease in GSIS (Fig. 2H). In insulin tolerance tests, MRS2905 also caused significant impairments in insulin sensitivity in lean and obese mice (Figs. 1J, 2I). MRS2905 treatment also led to significant increases in blood glucose levels in glucagon challenge tests in both lean and obese mice (Figs. 1K, 2J). As glucagon potently stimulates gluconeogenesis, we carried out a PTT to assess gluconeogenesis in vivo. Following MRS2905 and pyruvate treatment of obese mice, blood glucose levels were elevated 90 min post-injection, as compared with control mice treated with pyruvate alone (Figs.1L, 2K). Taken together, these data demonstrate that lean and obese mice display distinct impairments in whole body glucose homeostasis and plasma insulin levels after agonist-induced P2Y_14_R signaling.

### 3.2. Acute P2Y_14_R activation leads to the phosphorylation of JNK and upregulates hepatic gluconeogenic genes

To investigate the relationship between hyperglycemia and the JNK/G6PC pathway in the liver following MRS2905 treatment in lean and obese mice, JNK phosphorylation and the expression of gluconeogenic genes were examined. Liver tissues were collected 40 min after administration of vehicle or MRS2905 (10 mg/kg, i.p.) to lean and obese mice. Whole-liver lysates were used to assess c-Jun N-terminal kinase (JNK) phosphorylation, while the expression of gluconeogenic genes was analyzed at the transcript level. JNK phosphorylation was significantly increased in the liver lysates of both lean and obese mice 40 min after MRS2905 administration compared with vehicle-treated control mice (Figs. 3A, B, E, F). The hepatic transcript levels of the two key gluconeogenic genes, *G6pc* and *Pck1*, were also significantly upregulated in MRS2905-treated lean and obese mice (Figs. 3C, D, G, H). Uncropped and/or unprocessed images of the JNK and p-JNK blots are provided in Supplementary Fig. S4. Collectively, these findings suggest that activation of the JNK/G6PC pathway likely contributes to MRS2905-induced hyperglycemia. To confirm whether MRS2905-induced hyperglycemia is direct activation of G_i_-coupled P2Y_14_R. Glucose output assay carried out in primary hepatocytes isolated from HFD-consumed mice in presence or absence of MRS2905. Surprisingly, after treatment with MRS2905 (10 and 100 nM) did not show any alteration in glucose output whereas glucagon (100 nM) treatment increase glucose production from primary hepatocytes (Supplementary Fig. S2).

**Figure 3:**
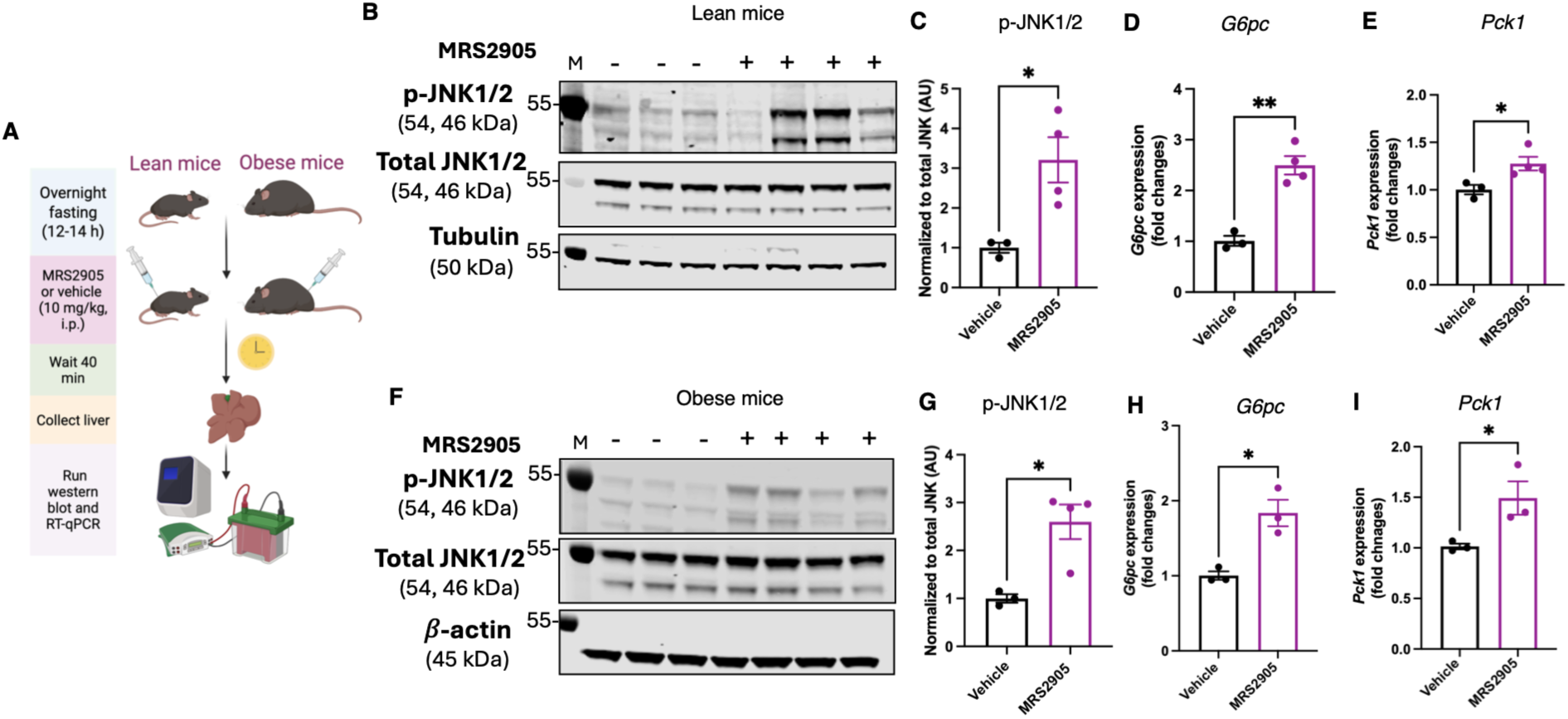
Acute activation of P2Y_14_R signaling upregulates hepatic JNK phosphorylation and gluconeogenic genes in lean and obese mice. **(A)** Schematic overview of acute MRS2905 treatment in lean and obese mice followed by tissue harvest and subsequent western blot and RT-qPCR analysis for JNK phosphorylation and gluconeogenic genes, respectively. **(B)** Immunoblot of liver lysates from lean mice received MRS2905 (10 mg/kg, i.p.). Liver tissue collected from lean mice 30 min after MRS2905 administered intraperitoneally and tissue lysate from saline and MRS2905 treated mice were subjected to Western blot. Total and phosphorylated JNK was detected using respective antibodies, n=3-4/group. **(C)** Quantification of the Western blotting data shown. Data are given as the means ±SEM. *P < 0.05, **P < 0.01, ***P < 0.001, and ****P < 0.0001, as compared with the corresponding saline treated control group [by 2-tailed unpaired Student’s t-test]. qRT-PCR analysis of rate-limiting gluconeogenic genes **(D)** G6pc, and **(E)** Pck1 mRNA levels in liver tissue obtained from lean mice 40 min after treated with saline or MRS2905 (n=3-4/group). **(F)** Immunoblot of liver lysates from mice maintained on obesogenic diet, received MRS2905 (10 mg/kg, i.p.). Liver tissue collected from obese mice 40 min after MRS2905 administered intraperitoneally and tissue lysates from saline and MRS2905 treated mice were subjected to Western blot. Total and phosphorylated JNK was detected using respective antibodies (n=3-4/group). **(G)** Quantification of the Western blotting data are given as the means ±SEM. *P < 0.05, **P < 0.01, ***P < 0.001, and ****P < 0.0001, as compared with the corresponding saline treated control group [[by 2-tailed unpaired Student’s t-test]. qRT-PCR analysis of two gluconeogenic genes **(H)** G6pc, and **(I)** Pck1 mRNA levels in liver tissue collected from obese mice 40 min after treated with saline or MRS 2905 (n=3-4/group). AU, arbitrary units.

### 3.3. Hyperglycemic effect of MRS2905 is partially blunted in adipo-P2ry14^-/-^ and abolished in WB-P2ry14^-/-^ mice

To assess whether MRS2905-induced hyperglycemia is attenuated in obese adipo-*P2ry14^-/-^* mice, MRS2905 or vehicle was acutely administered to both control and adipo-*P2ry14^-/-^* mice maintained on HFD for at least 8 weeks (Fig. 4A). In control mice, MRS2905 induced hyperglycemia compared with vehicle (Fig. 4B). However, in adipo-*P2ry14^-/-^* mice, the hyperglycemic response to MRS2905 was partially blunted relative to vehicle treatment (Fig. 4C). MRS2905 (10 mg/kg, i.p.) was administered to WB-*P2ry14^-/-^* mice are maintained on regular chow diet, and overnight fasting blood glucose levels was measured (Fig. 4D). WT control mice showed a robust hyperglycemic effect compared with vehicle treated mice (Fig. 4E). However, WB-*P2ry14^-/-^* mice showed no effect on fasting blood glucose (Fig. 4F). These data suggest that MRS2905 is selective for P2Y_14_R in mice, and the MRS2905-mediated hyperglycemic effect is completely P2Y_14_R-dependent, again emphasizing the selectivity of this agonist.

**Figure 4.**
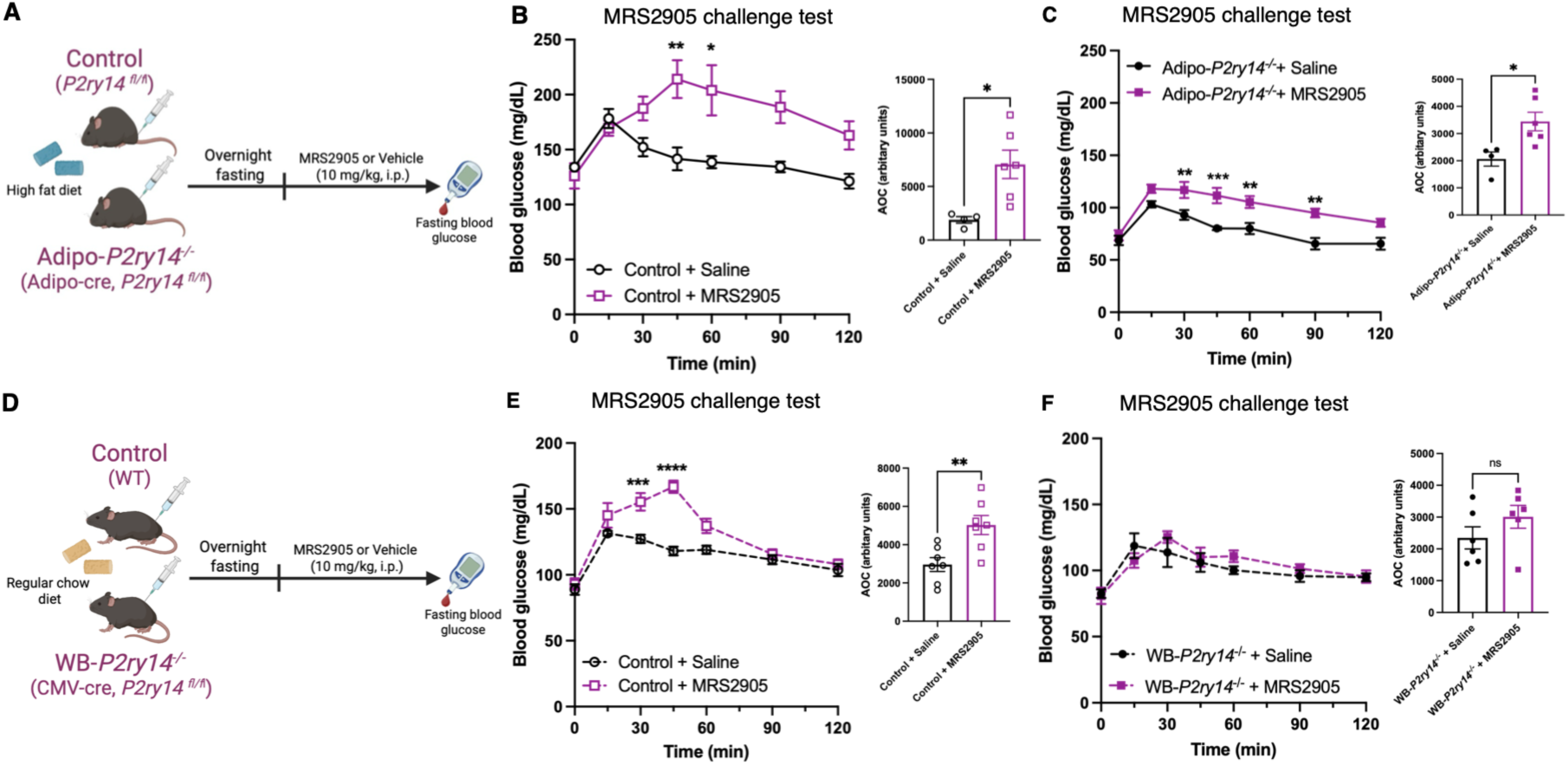
The hyperglycemic effect of MRS2905 is partially attenuated in HFD-fed adipo-P2Y_14_KO mice but completely abolished in lean WB-P2Y_14_KO mice. **(A)** Schematic presentation of control and adipo-P2ry14^-/-^ mice were maintained on HFD for at least 8 weeks and treated with saline or MRS2905 (10 mg/kg, i.p.). Effects of acute P2Y_14_R activation with MRS2905 in **(B)** control littermates and **(C)** adipocyte-specific *P2ry14* knockout (Adipo-*P2ry14*^⁻/⁻^) mice. **(D)** Schematic presentation of control and WB-P2ry14^-/-^ mice were maintained on a regular chow diet and treated with saline or MRS2905 (10 mg/kg, i.p.). Effects of acute P2Y_14_R activation with MRS2905 in **(E)** their age-matched control WT and **(F)** whole-body *P2ry14* knockout (WB-*P2ry14*^⁻/⁻^) mice.

### 3.4. Acute MRS4779 treatment improves whole-body glucose homeostasis in obese mice

To study the metabolic consequences of acute P2Y_14_R blockade in vivo, we administered the P2Y_14_R antagonist prodrug MRS4779 with two doses (10 mg/kg, i.p.) to WT obese mice, followed by a series of metabolic tests (Fig. 5A, B). Blocking P2Y_14_R by MRS4779 partially blunted the increased fasting hyperglycemia caused by MRS2905 (Fig. 5C). Two doses of MRS4779 were administered in obese mice and different metabolic studies conducted (Fig. 7D). A GTT showed that MRS4779 treatment greatly improved glucose tolerance (Fig. 7E). Moreover, GSIS was increased in MRS4779-treated mice (Fig. 7F). MRS4779 treatment of obese mice also resulted in improved glucose tolerance (Fig. 7G). These data suggest that acute treatment with a P2Y_14_R antagonist (mono-prodrug) improves metabolic health in obese mice.

**Figure 5.**
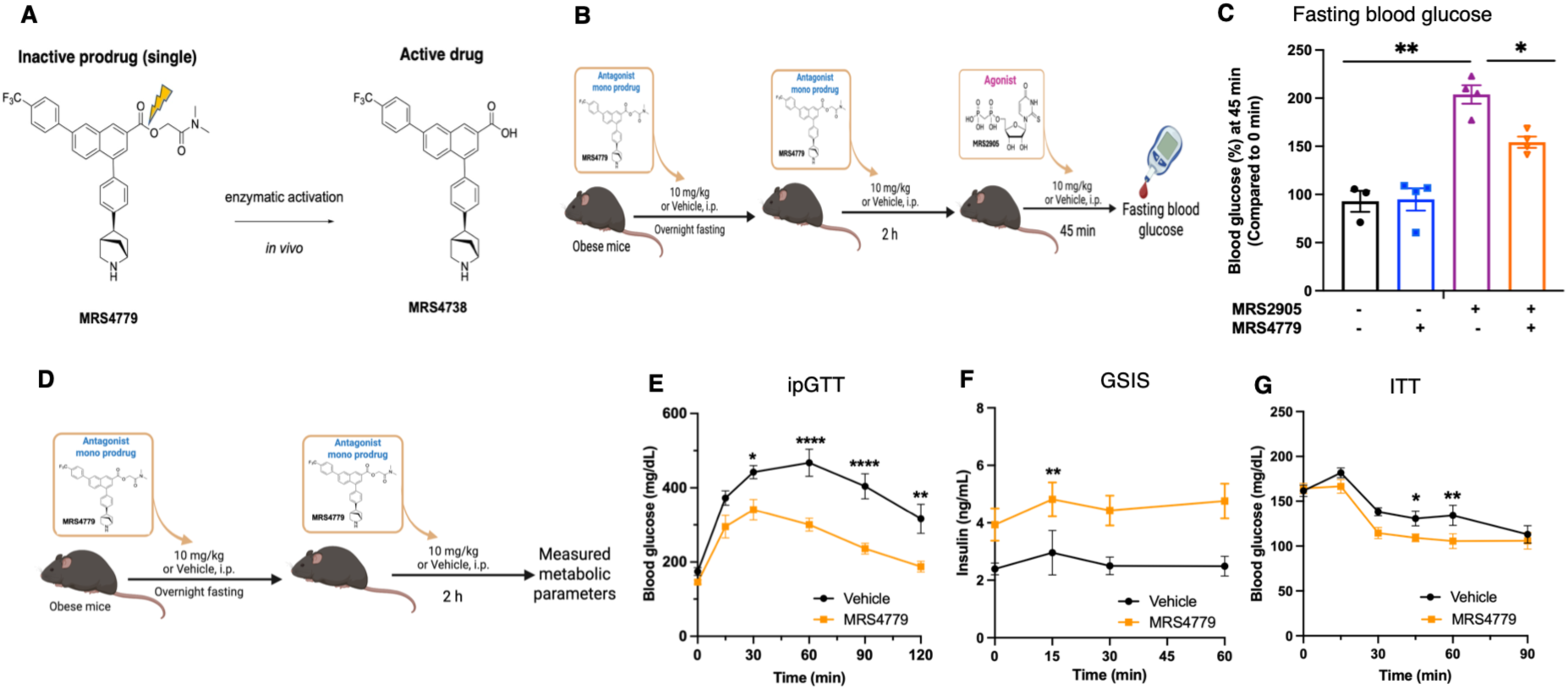
Acute pharmacological antagonism of P2Y_14_R improves glucose homeostasis in obese mice. **(A)** Chemical structures and schematic illustrating the conversion of the P2Y_14_R antagonist mono-prodrug MRS4779 to the active antagonist MRS4738. **(B)** Schematic presentation of the acute treatment protocol. Obese mice received vehicle or two doses of MRS4779 (10 mg/kg, i.p.), followed by vehicle or the P2Y_14_R agonist MRS2905 (10 mg/kg, i.p.), as indicated. Fasting blood glucose levels were measured after 45 min. **(D)** Schematic presentation of acute effect of MRS4779 in glucose homeostasis. To assess the effects of acute P2Y_14_R antagonism on systemic glucose homeostasis, obese mice received two doses of vehicle or MRS4779 (10 mg/kg, i.p.) before metabolic testing. **(C)** Intraperitoneal glucose tolerance (ipGTT), **(D)** glucose-stimulated insulin secretion (GSIS), and **(E)** insulin tolerance test (ITT) were performed as indicated. Data are presented as mean ± SEM (n=4–6 mice/group) for metabolic phenotyping. Statistical significance is indicated as *P < 0.05, **P < 0.001, ***P < 0.0001, and ****P < 0.00001.

### 3.5. Chronic MRS4779 treatment reduced body weight and improved whole-body glucose homeostasis in obese mice

We next evaluated the potential therapeutic efficacy of MRS4779 in obesity and T2D. Obese WT mice were chronically treated with MRS4779 at a dose of 5 mg/kg (i.p.) twice a day for 10 days. Prior to drug treatment, HFD-fed mice exhibited significantly greater body weight than chow-fed control mice (Fig. 6A). Over the treatment course, MRS4779 significantly reduced body weight in obese mice by ∼11% of total body weight compared to vehicle-treated obese mice (Fig. 6A, B). MRS4779 treatment did not affect lean mass (Fig. 6C) while it reduced fat mass and adiposity associated with obesity in DIO mice (Figs. 6D, E, F). We also found that chronic MRS4779 treatment lowered fed and fasting blood glucose levels (Figs. 6G, H). We monitored food intake in all mouse groups. MRS4779 reduced food intake during the first 3 days, and after the 4^th^ day food intake in the antagonist treated group returned to baseline (Fig. 6I). Surprisingly, chronic MRS4779 did not significantly affect glucose tolerance (Fig. 6J). Circulating triglyceride levels in antagonist-treated obese mice were not significantly different from vehicle-treated obese mice (Fig. 6K).

**Figure 6.**
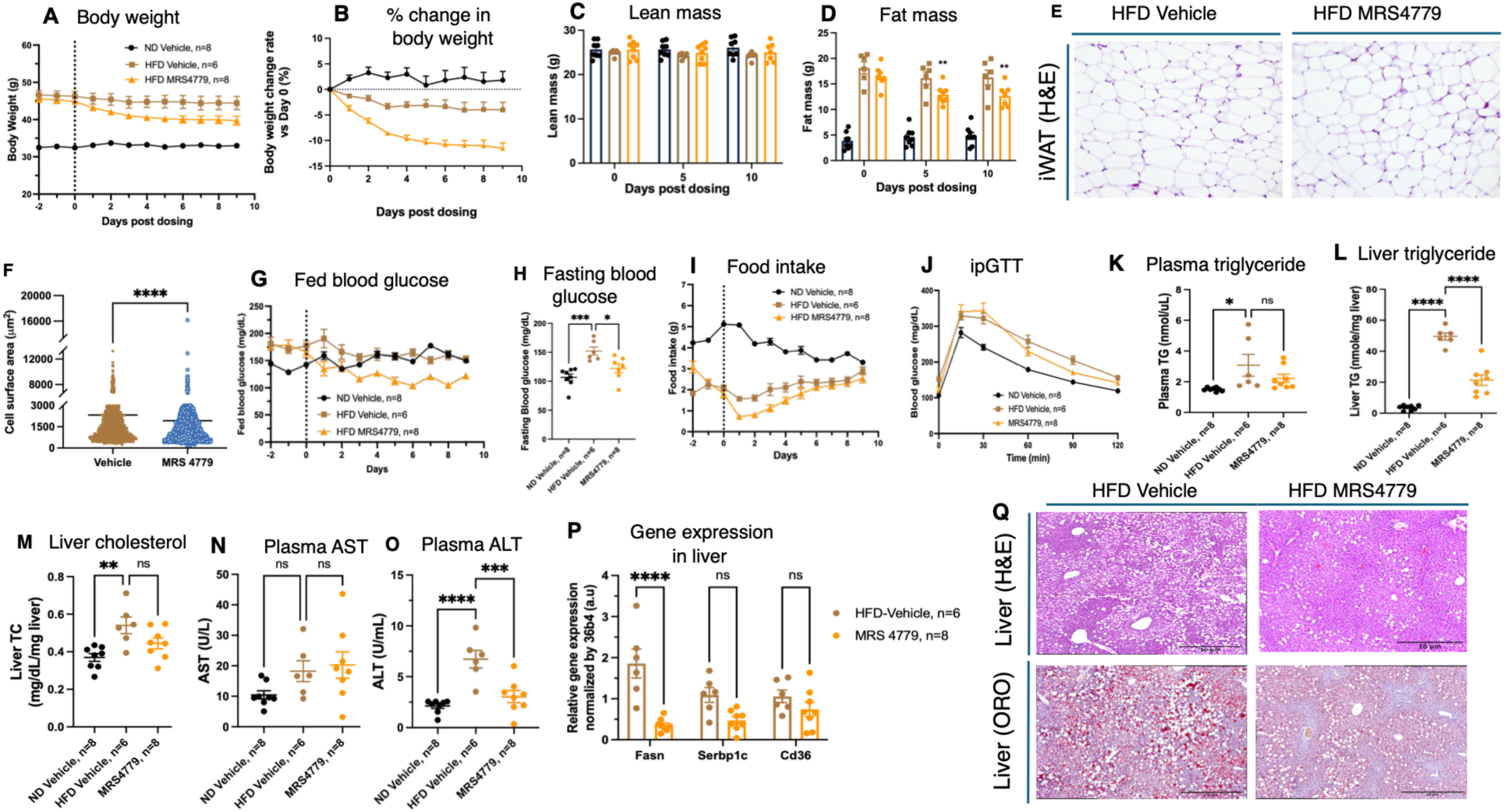
Chronic pharmacological antagonism of P2Y_14_R reduces body weight and improves glucose homeostasis in obese mice. Obese mice were treated with the P2Y_14_R antagonist prodrug MRS4779 twice daily for 10 days, and metabolic parameters were assessed as indicated. **(A)** Body weight, **(B)** percentage change in body weight, **(C)** lean mass, and **(D)** fat mass in mice maintained on a normal diet (ND) and treated with vehicle, or mice maintained on a high-fat diet (HFD) and treated with vehicle or MRS4779. **(E)** Representative H&E-stained sections of inguinal white adipose tissue (iWAT) and **(F)** quantification of adipocyte area in iWAT from vehicle- and MRS4779-treated obese mice. **(G)** Fed blood glucose levels measured at the indicated time points and **(H)** overnight-fasted blood glucose levels measured on day 12. **(I)** Cumulative food intake during the light and dark phases and **(J)** intraperitoneal glucose tolerance test (ipGTT). **(K)** Plasma triglyceride, **(L)** hepatic triglyceride, **(M)** hepatic cholesterol, **(N)** plasma aspartate aminotransferase (AST), and **(O)** plasma alanine aminotransferase (ALT) levels were measured in all three groups. **(P)** Hepatic mRNA expression of *Srebp1*, *Fasn*, and *Cd36* in vehicle- and MRS4779-treated obese mice. **(Q)** Representative H&E- and Oil Red O (ORO)-stained liver sections from vehicle- and MRS4779-treated obese mice. Data are presented as mean ± SEM (n = 6–8 mice/group). Statistical significance is indicated as *P < 0.05, **P < 0.001, ***P < 0.0001, and ****P < 0.00001.

Obesity increases the risk of developing MASLD, a condition marked by hepatic steatosis caused by an imbalance in hepatic fatty acid uptake, synthesis, oxidation, and export [28]. Given the prominent effects of MRS4779 on body weight, fasting blood glucose, and adiposity, we then investigated its impact on liver steatosis. We found that MRS4779 treatment reduced liver triglyceride level significantly without notably changing liver cholesterol level (Figs. 6L, M). Plasma AST remained unchanged in the MRS4779-treated group compared to vehicle-treated obese mice, but ALT was lowered in the antagonist-treated group (Figs. 6N, O). Moreover, MRS4779 treatment improved hepatic steatosis, as determined by H&E staining of liver sections (Fig. 6Q). This improvement was further supported by a decrease in ORO staining in MRS4779-treated liver sections compared to the HFD vehicle-treated control group (Fig. 6Q). To investigate the molecular mechanism underlying the effect of MRS4779 on hepatic fat accumulation, we evaluated the expression of key genes involved in hepatic lipid metabolism using qRT-PCR in DIO mice with or without MRS4779 treatment. MRS4779 administration significantly downregulated fatty acid synthase (*Fasn*), sterol regulatory element-binding protein 1 (*Srebp1*) and fatty acid transporter (*Cd36*) transcript levels (Fig. 6P). Overall, these findings suggest that MRS4779 has a beneficial effect on hepatic lipid accumulation and liver function in the context of obesity-associated MASLD.

### 3.6. Hepatocyte-specific P2Y_14_ R KO mice show reduced fasting blood glucose but no effect on other metabolic parameters

To determine the metabolic roles of hepatic *P2ry14*, we first quantified *P2ry14* mRNA levels in whole liver from lean and obese mice. Interestingly, consumption of HFD increased hepatic *P2ry14* levels (Fig. 7A). We obtained the same result with isolated hepatocytes prepared from lean and obese mice (Fig. 7B). To explore the physiological role of P2Y_14_R in hepatocytes, we generated mice lacking P2Y_14_R specifically in hepatocytes (Hep-*P2ry14^-/-^*mice) (Fig. 7C). Male Hep-*P2ry14^-/-^*mice and their control littermates (*P2ry14^fl/fl^* mice) were maintained on RC. On RC, Hep-*P2ry14^-/-^* and control mice (males) did not exhibit any significant differences in fed and fasting blood glucose, plasma insulin, FFA, triglyceride, cholesterol, glucose tolerance, as well as insulin and glucagon sensitivity (Supplementary Figs. S5 A-H). We obtained similar results with female Hep-*P2ry14^-/-^* mice and their control littermates fed RC diet (Supplementary Figs. S6 A-H).

**Figure 7.**
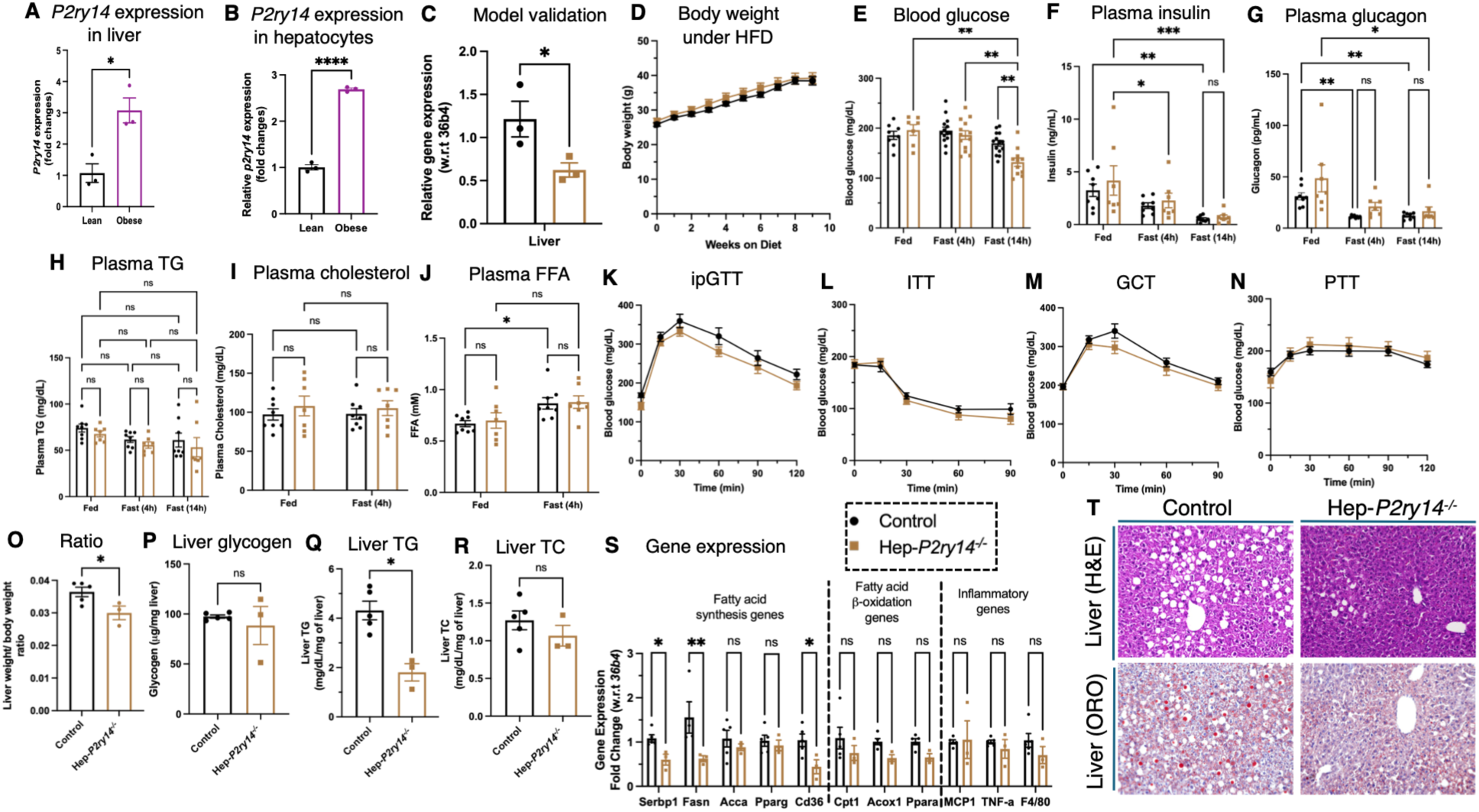
Hepatic P2Y_14_R expression is altered in obesity, and hepatocyte-specific P2Y_14_R deletion improves fasting glucose levels in male mice maintained on a high-fat diet. Relative *P2ry14* mRNA expression levels in **(A)** liver and **(B)** primary hepatocytes isolated from lean and obese WT C57BL/6NTac mice (n = 3/group). **(C)** Hepatic *P2ry14* mRNA expression in Hep-*P2ry14*^⁻/⁻^ mice and control littermates, confirming the loss of *P2ry14* expression in the liver (n = 3/group). Metabolic parameters were assessed in male Hep-*P2ry14*^⁻/⁻^ mice and control littermates maintained on an obesogenic diet for at least 8 weeks. **(D)** Body weight, **(E)** blood glucose, **(F)** plasma insulin, and **(G)** glucagon levels were measured under fed, 4-h-fasted, and 14-h-fasted conditions. Plasma **(H)** triglyceride, **(I)** cholesterol, and **(J)** free fatty acid (FFA) levels were measured under fed and 4-h-fasted conditions. **(K)** Intraperitoneal glucose tolerance test (IPGTT), **(L)** insulin tolerance test (ITT), **(M)** glucagon challenge test (GCT), and **(N)** pyruvate tolerance test (PTT) performed in Hep-*P2ry14*^⁻/⁻^ mice and control littermates. **(O)** Ratio of liver weight and body weight of individual mice is presented (n=3-5/group). Liver **(P)** glycogen, **(Q)** total triglyceride, **(R)** total cholesterol measured in knock out and control group of mice (n=3-5/group). **(S)** Genes related to fatty acid synthesis, fatty acid beta oxidation and inflammatory genes quantified using RT-qPCR (n=3-5/group). **(T)** Representative H&E- and Oil Red O (ORO)-stained liver sections from control and Hep-*P2ry14*^⁻/⁻^mice. Data are presented as mean ± SEM (n = 7–14 mice/group for metabolic phenotyping). Statistical significance is indicated as *P < 0.05, **P < 0.001, ***P < 0.0001, and ****P < 0.00001.

Next, we maintained male Hep-*P2ry14^-/-^* mice and their control littermates on an HFD for 8 weeks and measured body weight weekly. Hep-*P2ry14^-/-^* and control mice show comparable increases in body weight (Fig. 7D). However, after an overnight fast, blood glucose levels were significantly lower in Hep-*P2ry14^-/-^* mice (Fig. 7E). Fasting plasma insulin and glucagon levels, insulin sensitivity, and pyruvate tolerance were similar between the two groups of mice (Figs. 7F, G, L, N). Glucose- and glucagon-induced elevations in blood glucose levels were not significantly altered in Hep-*P2ry14^-/-^* mice compared with HFD control littermates (Figs. 7K, M). Fed and fasting plasma cholesterol and FFA levels remained unchanged (Figs. 7I, J). Ratio of liver weight and body weight was significantly reduced in knockout mice compared to control mice whereas liver glycogen level remains unaltered (Figs. 7O, P). Liver total triglyceride level is significantly decrease without change of total cholesterol levels in Hep-*P2ry14^-/-^* mice compared with control (Figs. 7Q, R). Genes related to fatty acid synthesis (*serbp1*, f*asn* and *cd36*) are decreased in knock out mice compared to control mice without change of fatty acid oxidation and inflammatory genes (Fig. 7S). Moreover, hepatic steatosis was determined by H&E staining of liver sections (Fig. 7T). This improvement was further supported by a decrease in ORO staining in liver sections of Hep-*P2ry14^-/-^*control group (Fig. 7T) These findings suggest that lack of P2Y_14_R on hepatocytes did not impact glucose homeostasis in the mice maintained on HFD but decreased hepatic lipid accumulation by reducing liver steatosis in HFD-induced obese mice.

### 3.7. P2Y_14_R antagonists improve lipid accumulation in an in vitro steatosis model

We selected mouse AML12 cells to study effects of P2Y_14_R antagonists in the palmitic acid (PA)-induced in vitro steatosis model. We first measured the mRNA expression of *P2ry14* in PA-treated AML12 cells after 24, 48 and 72 h. *P2ry14* expression was increased time-dependently (Fig. 8A), and 48 h was selected for further studies. We utilized two selective and potent antagonists, PPTN and MRS4738 which is a mono-prodrug form of MRS4779. Neither PPTN nor MRS4738 at 10 µM showed any cytotoxicity in AML12 mouse hepatocyte cells (Fig. 8B). After 48 h of PA treatment, lipid accumulation was detected by ORO staining, and PA treatment increased lipid droplet numbers compared to control cells (Fig. 10C). When PPTN or MRS4738 was co-treated with PA for 48 h, both antagonists reduced lipid droplet accumulation in AML12 hepatocytes, at 400 and 500 nM, respectively, compared to cells treated with PA alone. However, at the lowest concentration (100 nM), neither antagonist showed any prominent changes in lipid accumulation compared to only PA treated cells (Fig. 8C).

**Figure 8.**
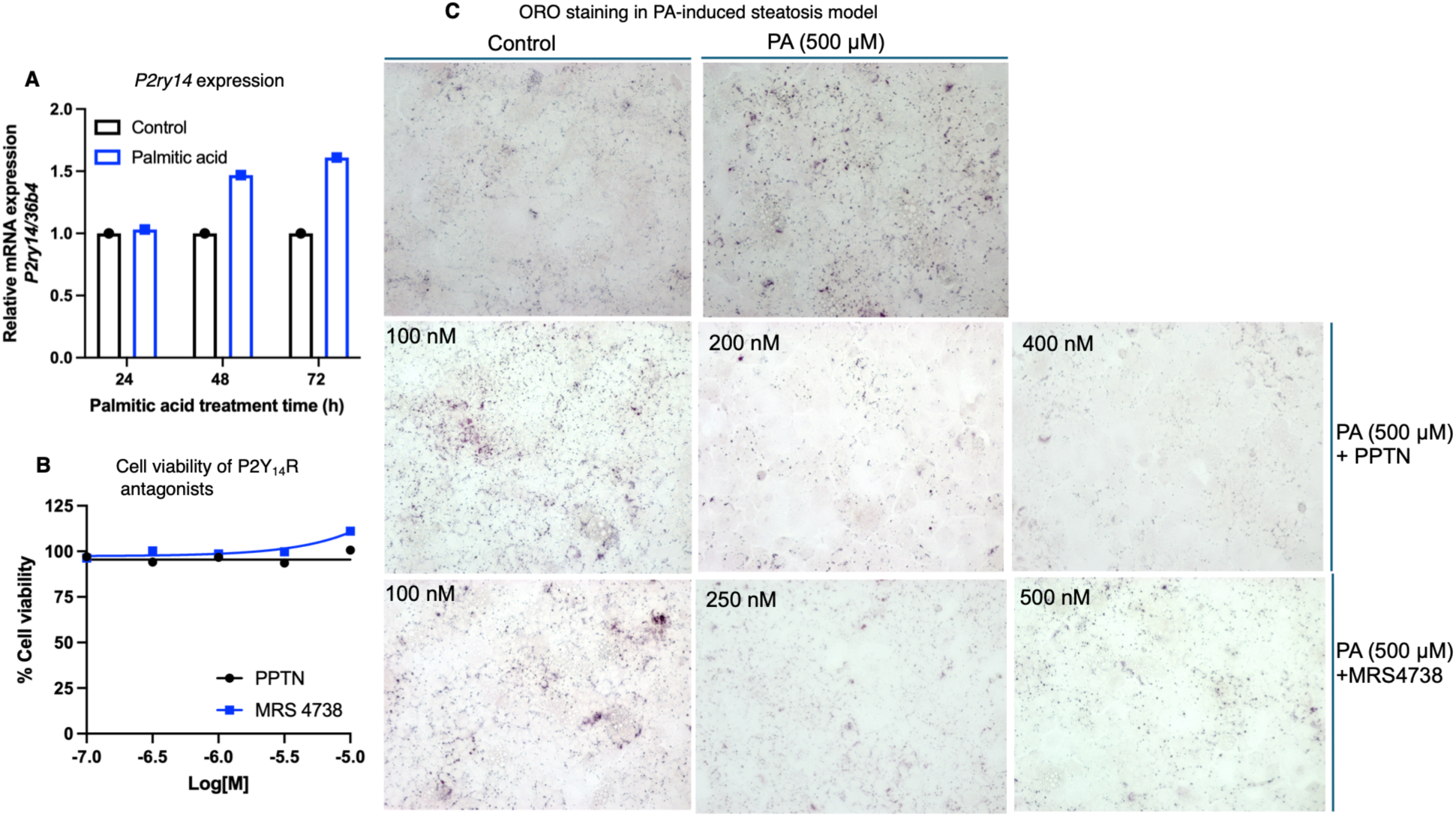
P2Y_14_R antagonists reduce palmitic acid-induced lipid accumulation in an in vitro model of hepatic steatosis. **(A)** AML12 mouse hepatocytes were treated with palmitic acid (PA) to induce an in vitro steatosis model. Cells were treated with PA for 24, 48, or 72 h, followed by assessment of *P2ry14* mRNA expression. **(B)** Cell viability following treatment with the P2Y_14_R antagonists PPTN and MRS4738 was assessed in AML12 cells using an MTT assay. **(C)** Intracellular lipid accumulation was evaluated by Oil Red O (ORO) staining following co-treatment of AML12 cells with PA (500 µM) and different concentrations of the P2Y_14_R antagonists.

### 3.8. Hepatic P2Y_14_R expression does not correlate with metabolic dysfunction in humans

The *P2RY14* gene is expressed in human adipose tissue at moderate levels [11]. Its expression is altered in the subcutaneous fat of obese individuals with insulin resistance compared to matched healthy individuals [14]. In contrast, healthy human liver tissue exhibits low levels of *P2RY14* expression [11]. This observation encouraged us to investigate whether metabolic dysfunction influences hepatic *P2RY14* mRNA expression in individuals with metabolic dysfunction–associated steatotic liver disease (MASLD). *P2RY14* gene expression was quantified from subcutaneous adipose tissues of overnight fasted human subjects with MASLD (n=29). Adipose expression of *P2RY14* was negatively associated with fasting glucose (p=0.052; R=-0.37, Supplementary Fig. S7A), with a trending positive correlation to HOMA-B (p=0.11; R =0.31, Supplementary Fig. S7B). There was no association between adipose *P2RY14* expression and the area under the GTT curve (Supplementary Fig. S7C). These data suggest that alteration of *P2RY14* expression in adipose tissue is specific to the modulation of fasting blood glucose levels. There was also no association between adipose *P2RY14* expression and hepatic steatosis (Supplementary Fig. S7D). Notably, *P2RY14* mRNA was not detected in liver tissue from the same individuals.

## 4. Discussion

P2Y_14_R has emerged as a promising therapeutic target in several chronic and acute inflammatory disorders, including chronic neuropathic pain, asthma, inflammatory bowel disease (IBD), and acute kidney injury (AKI) [19, 29–31]. More recently, P2Y_14_R has also been proposed as a potential metabolic target for obesity and type 2 diabetes [12–14]. In the present study, we systematically investigated the metabolic functions of P2Y_14_R using complementary pharmacological (agonists and antagonists) and genetic (P2Y_14_R knockout mouse models) approaches. We characterized both endogenous and synthetic agonists to identify a suitable pharmacological agonist for selectively activating P2Y_14_R in vivo with high selectivity and metabolically stable. Subsequently examined the metabolic consequences of acute P2Y_14_R activation in lean and obese mice. Surprisingly, MRS2905 showed high selectivity with metabolic stability in mice, and acute P2Y_14_R activation impaired whole body glucose metabolism in lean and DIO mice. In contrast, acute blockade of P2Y_14_R with MRS4779 partially blunted MRS2905-induced hyperglycemia in obese mice indicative of the selectivity of this agonist in mice. Additionally, chronic treatment with mono-prodrug antagonist MRS4779 led to a body weight reduction and improved liver steatosis but had no effect on glucose homeostasis. *P2Y_14_R* gene expression was upregulated in liver and hepatocytes from mice on HFD. Surprisingly, P2Y_14_R deletion from hepatocytes did not alter glucose metabolism; however, it improved lipid metabolism and reduced lipid accumulation.

UDP-glucose (UDP-G) is an intracellular pyrimidine nucleotide sugar that serves as a key intermediate in multiple metabolic pathways, including glycogen biosynthesis, glycolysis, and galactose metabolism [32]. Under conditions of cellular stress or tissue injury, however, UDP-glucose can be released into the extracellular space, where it functions as a damage-associated molecular pattern (DAMP) and activates P2Y_14_R through autocrine or paracrine signaling [33, 34]. Although UDP-glucose is the endogenous ligand for P2Y_14_R, several pharmacological limitations restrict its utility for in vivo studies. UDP-glucose is rapidly degraded by extracellular ectonucleotidases particularly in the liver or ectonucleotide pyrophosphatase/phosphodiesterase (ENPP5), resulting in a short duration of action following intraperitoneal or intravenous administration in mice [35–37]. Furthermore, at higher concentrations, UDP-glucose lacks receptor selectivity and can activate other purinergic receptors, including P2Y_2_R and P2Y_6_R [17]. Previous studies have therefore relied on relatively high doses of UDP-glucose for in vivo study. For example, Bassil et al. [38] administered 2,000 mg/kg UDP-glucose intraperitoneally to investigate the role of P2Y_14_R in gastric emptying. In addition, UDP-glucose has P2Y_14_R-independent biological activities. It functions as a molecular glue that promotes cereblon-mediated glucokinase degradation and alters insulin secretion [39]. Chronic administration of UDP-glucose (8 mg/kg, i.p.) has also been reported to improve hepatic steatosis and promote glycogenesis in diet-induced obese mice by inhibiting site-1 protease (S1P)-mediated cleavage of sterol regulatory element-binding proteins (SREBPs) [40]. Together, these studies underscore the pleiotropic actions of UDP-glucose and highlight the difficulty of attributing its in vivo metabolic effects exclusively to P2Y_14_R activation. Consequently, selective pharmacological agonists are required to accurately define the physiological and metabolic functions of P2Y_14_R.

The present study provides, to our knowledge, the first comprehensive characterization of the acute metabolic effects of selective P2Y_14_R agonists in a lean and obese mouse model. Synthetic agonists such as MRS2690 and MRS2905, (Supplementary Figure S8), which are UDP and UDP-G analogues, respectively, were developed to overcome the pharmacokinetic and selectivity limitations of UDP-glucose and UDP by exhibiting greater receptor selectivity and improved metabolic stability [17]. Consistent with these properties, our pharmacokinetic analysis demonstrated that UDP-glucose displayed a very short plasma half-life following a single intraperitoneal injection (10 mg/kg), with detectable plasma concentrations lasting less than 15 minutes and complete clearance by approximately 30 minutes, likely because of rapid hepatic metabolism and enzymatic degradation. In contrast, MRS2905 exhibited substantially greater metabolic stability and maintained bioactive plasma concentrations for at least 30 minutes after intraperitoneal administration, making it a more suitable agonist for in vivo P2Y_14_R activation. Thus, pharmacokinetic data suggested that prolonged in vivo stability of MRS2905 may be attributed to the methylene bridge within its phosphate moiety (Supplementary Figure S8), which confers greater resistance to nucleotidase-mediated degradation compared with a conventional diphosphate group in UDP or UDP-G.

Using these pharmacological tools, we compared the metabolic effects of three P2Y_14_R agonists (UDP-glucose, MRS2905, and MRS2690) in obese mice. Among these compounds, MRS2905 produced the greatest impairment in glucose metabolism, whereas UDP-glucose elicited only modest effects. MRS2690 induced a moderate elevation in fasting blood glucose, consistent with its lower potency relative to MRS2905. EC_50_ values in cAMP functional assay for MRS2690 and MRS2905 are 49 nM and 0.92 nM, respectively [17]. Thus, our in vivo efficacy study for these two synthetic agonists is also corroborate with previous structure activity relationships (SAR) studies for agonists on P2Y_14_R. These findings establish MRS2905 as a robust pharmacological probe for studying P2Y_14_R function in vivo. Importantly, acute activation of P2Y_14_R with MRS2905 impaired glucose homeostasis in both lean and DIO mice, providing direct evidence that selective P2Y_14_R activation acutely disrupts metabolic regulation. Collectively, these results identify MRS2905 as an effective in vivo tool for investigating P2Y_14_R biology and further support P2Y_14_R as a potential therapeutic target in metabolic disease.

Previous studies have shown that activation of hepatic G_i_-coupled GPCR signaling enhances JNK activity, leading to increased hepatic glucose production (HGP) and hyperglycemia in G_i_-DREADD mice [21]. Activation of hepatic JNK signaling also promotes the expression of key gluconeogenic genes, thereby facilitating glucose production by the liver [41]. Acute P2Y_14_R activation with MRS2905 increased hepatic JNK phosphorylation and upregulated gluconeogenic gene expression (*G6pc*, *Pck1*) in lean and obese mice. However, the activation of P2Y_14_R with MRS2905 in mouse primary hepatocytes from obese mice didn’t show any alteration in glucose output. Mechanistically, acute P2Y_14_R activation with MRS2905 increased fasting blood glucose level, elevate circulating glucagon levels, reduce plasma insulin, enhance hepatic JNK phosphorylation, and upregulate the expression of key gluconeogenic genes. These observations suggest that P2Y_14_R activation disrupts endocrine regulation of glucose homeostasis. One possible mechanism is that MRS2905 directly activates P2Y_14_R expressed in pancreatic islets, suppressing insulin secretion from β-cells while stimulating glucagon release from α-cells as a counter-regulatory response secondary to reduced insulin secretion and the resulting hyperglycemia. Regardless of the initiating mechanism, reduced insulin levels would impair peripheral glucose disposal, whereas elevated glucagon would promote hepatic glucose production under fasting and fed conditions, together contributing to the transient hyperglycemia in fed and fasted mice observed following MRS2905 administration.

Interestingly, despite increased hepatic gluconeogenic gene expression in vivo, MRS2905 failed to stimulate glucose production in isolated primary hepatocytes. This finding suggests that the hyperglycemic response is unlikely to result from a not the direct effect of P2Y_14_R activation in hepatocytes. Instead, the induction of hepatic gluconeogenic pathways in vivo is more likely secondary to systemic hormonal changes, particularly reduced insulin and elevated glucagon levels, or to indirect signaling through extrahepatic tissues. These results support a model in which P2Y_14_R regulates glucose homeostasis primarily through endocrine and inter-organ communication rather than through direct modulation of hepatocyte glucose production.

In contrast, acute treatment of mice with a P2Y_14_R antagonist (mono-prodrug MRS4779) resulted in lowered fasting plasma insulin levels and reversed the agonist-induced metabolic changes, thus restoring whole body glucose homeostasis. Cannabinoid receptor type-1 (CB1) is a G_i/o_-coupled receptor that is considered as a potential target for anti-obesity therapy [42]. Peripherally restricted pharmacological blockade of CB1 receptors with JD5037 reduces obesity-linked phenotypes such as body weight, fat mass and cardiometabolic risk, and improves hepatic steatosis in DIO mice [43, 44]. P2Y_14_R is also a G_i/o_-coupled receptor and chronic blockade of this receptor in DIO mice reduced body weight and fat mass. The beneficial effect on liver steatosis observed after MRS4779 administration suggests that P2Y_14_R antagonists may become clinically useful for the treatment of fatty liver. Thus, drug-like P2Y_14_R antagonists, such as MRS4779 and related analogues, may have potential for the treatment of obesity and obesity-related metabolic disorders.

## 5. Limitation of the study

Several limitations and important questions remain unresolved in the present study. The efficacy of standard antagonist PPTN or MRS4738 derived from mono-prodrug (MRS4779) as a standalone treatment was not evaluated in obese mice, either upon acute or chronic administration. Although the prototypical antagonists have low oral bioavailability and/or a hydrophobic nature that restricts achieving full efficacy in vivo, they are highly selective for P2Y_14_R with sub-nanomolar potency. This study provides a rational basis for the development of orally bioavailable P2Y_14_R antagonists for the treatment of obesity and T2D. Although P2Y_14_R is expressed in pancreatic islets, its specific roles in different islet cell populations remain unclear. A detailed investigation of how peripheral administration of P2Y_14_R-selective agonists affects insulin and glucagon secretion through distinct cell types would provide valuable mechanistic insights. P2Y_14_R expression in human hepatocytes has not been validated in metabolic diseases such as obesity, type 2 diabetes, and metabolic MASLD. Here, we cannot provide a detailed mechanism underlying the reduction of hepatic lipid accumulation upon antagonist treatment. Demonstrating P2Y_14_R expression in these clinically relevant conditions would enhance the translational significance of the findings and strengthen the rationale for targeting P2Y_14_R and development of antagonists as a therapeutic strategy for metabolic disorders.

## Acknowledgements

We thank the NIDDK Intramural Research Program for support (ZIA DK031116; ZIA DK031129). We thank Mouse Metabolism Core (NIDDK; 1ZICDK070002) for carrying out several measurements and advice, Clinical Mass Spectrometry Core (NIDDK), Dr. Jeffrey Reece (Advanced Light Microscopy & Image Analysis Core, NIDDK) provided helpful advice regarding the imaging. The contributions of the NIH authors are considered Workers of the United States Government. The findings and conclusions presented in this paper are those of the authors and do not necessarily reflect the views of the NIH or the U.S. Department of Health and Human Services. BioRender (https://www.biorender.com/) was used to draw schematic presentations and graphical abstract.

## Author Contributions

Conceived experimental design: AP, KAJ; Performed experiments: AP, HC, PV, PJW, TD, OG, SP, HS, RU, AW; Contributed research materials: SHK, ZW; Analysis of data and writing of manuscript: AP, PV, PJW, OG, AW, YR, JW, KAJ; Wrote first draft: AP. All authors reviewed and approved the final manuscript version.

## Data and materials availability

All data needed to evaluate the conclusions in the paper are present in the paper and/or the Supplementary Materials. Additional data related to this paper may be requested from the authors.

## Abbreviations

DIO: diet induced obesity
G6pc: glucose-6-phosphatase, catalytic subunit
Pck1: phosphoenolpyruvate carboxykinase
H&E: Hematoxylin and Eosin
HFD: high fat diet
HSC: hepatic stellate cell
ORO: Oil Red O
PA: palmitic acid
AML12: Alpha mouse liver 12
GPCR: G protein-coupled receptor
RCD: regular chow diet
T2D: Type 2 diabetes
UDPG/ UDP-glucose: uridine-5′-diphosphoglucose
UDP: uridine 5′-diphosphate.

