## Supplementary Information for "P2Y14 receptor agonist impairs, and antagonist improves whole-body glucose homeostasis and liver steatosis"

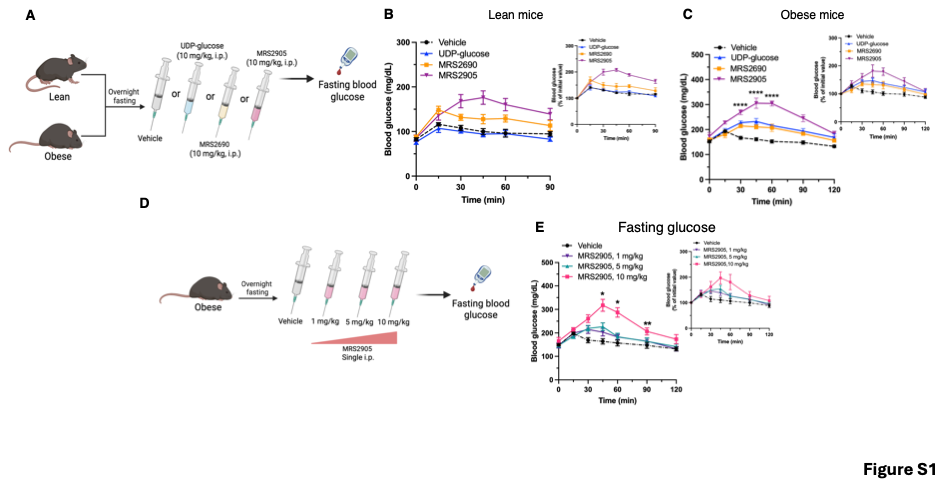

### **Figure S1. Acute effects of P2Y_14_ receptor ligands on fasting blood glucose in lean and obese mice** (A) Schematic of the experimental design. Lean and obese mice were fasted overnight and administered vehicle, UDP-glucose (10 mg/kg, i.p.), MRS2690 (10 mg/kg, i.p.), or MRS2905 (10 mg/kg, i.p.), followed by measurement of blood glucose over time. (B, C) Blood glucose concentrations and corresponding changes relative to initial glucose levels following administration of the indicated compounds in lean (B) and obese (C) mice. (D) Schematic of the dose-response experiment in overnight-fasted obese mice administered vehicle or MRS2905 (1, 5, or 10 mg/kg, single i.p. injection). (E) Blood glucose concentrations and corresponding changes relative to initial glucose levels following administration of increasing doses of MRS2905. Data are presented as shown in the figure. *P < 0.05 and ****P < 0.0001, as indicated.

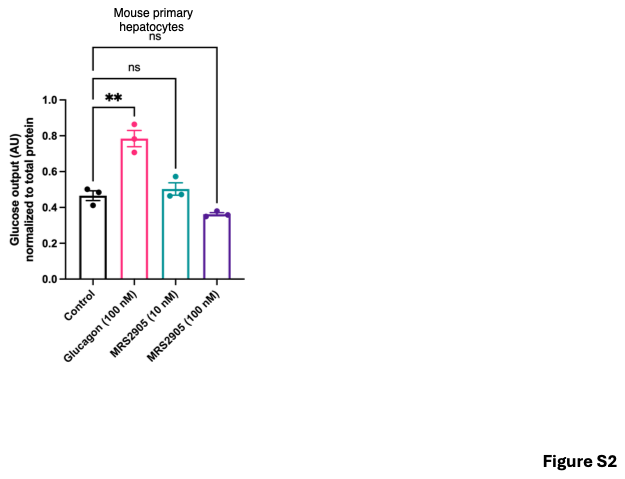

**Figure S2. Effects of P2Y_14_ receptor agonists on glucose output from primary mouse hepatocytes.** Glucose output from primary mouse hepatocytes isolated from obese mice treated with vehicle (control), glucagon (100 nM), MRS2905 (10 nM), or MRS2905 (100 nM). Glucose output is expressed in arbitrary units (AU) normalized to total protein. Individual data points and summary values are shown. **P < 0.01; ns, not significant, for the indicated comparisons.

**Supplementary methods**

**UPLC-MS/MS analysis of UDP-galactose, UDP-glucose, and MRS2905:**

High-performance liquid chromatography (HPLC) grade solvents, modifiers and standards were purchased from Sigma-Aldrich (St. Louis, MO, USA). Detection and quantification were performed using ultra-performance liquid chromatography–tandem mass spectrometry (UPLC-MS/MS) with a Thermo Scientific Vanquish UPLC system coupled to a Thermo Scientific Altis triple quadrupole mass spectrometer equipped with heated electrospray ionization (HESI-II) in negative ion mode at 4000 V.

UDP-glucose, UDP-galactose and MRS2905 calibration stock solutions (10–250 ng/mL) were prepared in water and stored at 4 °C. The internal standard solution, UDP-¹³C₆-glucose (50 ng/mL in water), was stored at −80 °C.

For sample preparation, 5 µL of calibration stock or plasma sample was mixed with 5 µL internal standard solution and 40 µL acetonitrile (ACN) in a 1.7 mL centrifuge tube, vortexed for 10 minutes, then centrifuged at 14,000 rpm for 15 minutes at 4 °C. The supernatant was transferred to an LC-MS vial, dried at room temperature under nitrogen (N₂), and reconstituted in 20 µL mixture of mobile phase A and mobile phase B (95:5, v/v), vortexed for 10 minutes prior to LC-MS analysis.

Chromatographic separation for UDP-glucose and UDP-galactose was achieved using a Thermo Scientific Hypercarb column (2.1 × 100 mm, 3 µm) maintained at 40 °C. Mobile phase A consisted of water containing 0.1% formic acid (FA), adjusted to pH 9 with ammonium hydroxide (NH₄OH), and mobile phase B was acetonitrile. The injection volume was 5 µL, and the flow rate was 125 µL/min. The gradient began at 5% B and was held until 1.5 min, increased to 20% B at 8 min, then rapidly increased to 95% B at 8.1 min, maintained until 11 min, and returned to 5% B at 11.1 min, for a total run time of 15 min. All samples were analyzed in duplicate.

Chromatographic separation for MRS 2905 was achieved using a Waters Acquity UPLC BEH C18 column (2.1 × 100 mm, 1.7 µm) maintained at 40 °C. Mobile phase A consisted of water containing 0.1% triethylamine (TEA), adjusted to pH 9 with formic acid (FA), and mobile phase B was acetonitrile. The injection volume was 2 µL, and the flow rate was 300 µL/min. The gradient initiated at 5% B, held until 3 min, increased to 90% B at 3.5 min, maintained until 5.5 min, and returned to starting conditions at 6 min, for a total run time of 8.5 min. All samples were analyzed in duplicate.

Quantitation of UDP-glucose, UDP-galactose, and UDP-¹³C₆-glucose (Internal Standard) was based on their multiple reaction monitoring (MRM) transitions and retention times. The transition *m/z* 565.09 → 323.13 was used for both UDP-glucose and UDP-galactose, while *m/z* 571.212 → 323.155 was used for UDP-¹³C₆-glucose. UDP-glucose and UDP-galactose retention time is 5.98 and 5.78 respectively. The calibration curve exhibited a minimum R^2^ ≥ 0.99 with 1/x weighting, meeting the FDA LC-MS guideline for linearity and quantitation.

Quantitation of MRS2905 and UDP-^13^C_6_-Glucose were based on multiple reaction monitoring (MRM) transitions *m/z*, 417.038 → 156.917 and 271.071 for MRS 2905, 571.212→ 323.155 for UDP-^13^C_6_-Glucose, respectively. The calibration curve exhibited a minimum R^2^ ≥ 0.99 with 1/x weighting, meeting the FDA LC-MS guideline for linearity and quantitation.

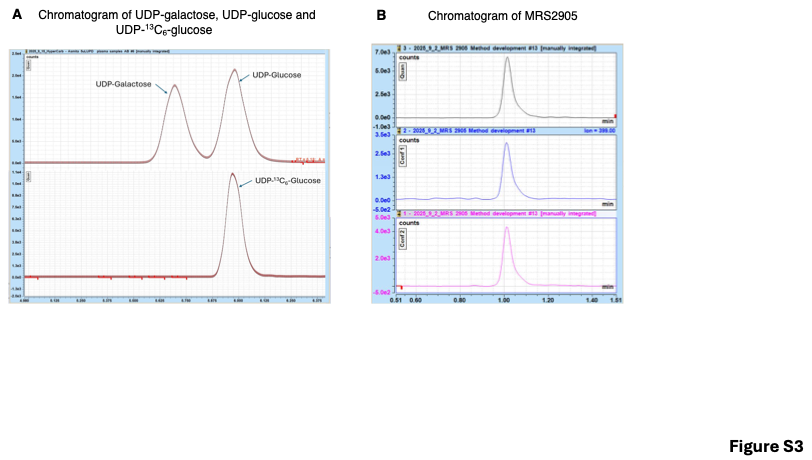

**Figure S3. Representative UPLC–MS/MS chromatograms of UDP-sugars, MRS2905, and internal standard.** Representative chromatogram of **(A)** UDP-galactose, UDP-glucose, and UDP-^13^C_6_-glucose obtained by ultra-performance liquid chromatography–tandem mass spectrometry (UPLC–MS/MS). **(B)** Representative chromatogram of MRS2905 monitored using three multiple-reaction monitoring (MRM) transitions. Peak retention time and quantification of analytes were confirmed using UDP-^13^C_6_-glucose as an internal standard.

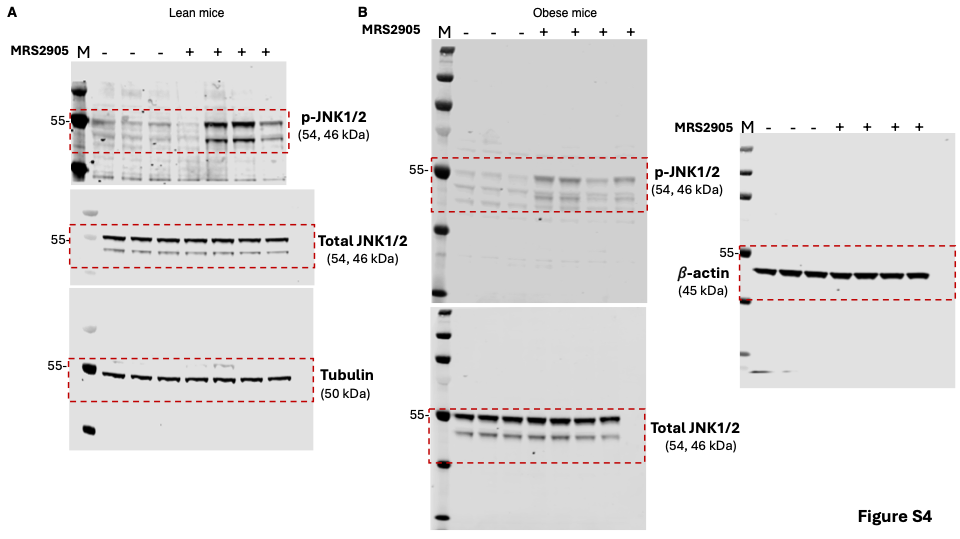

**Figure S4. Uncropped immunoblot images.** Nitrocellulose membrane images corresponding to the immunoblots of total JNK1/2, pJNK1/2 and loading control (tubulin or 𝛽-actin) shown in Figure 3B and 3F are provided here. All lanes used for the corresponding band intensity analysis and molecular weight markers are shown. The regions displayed in the main figures are indicated by red doted rectangular boxes.

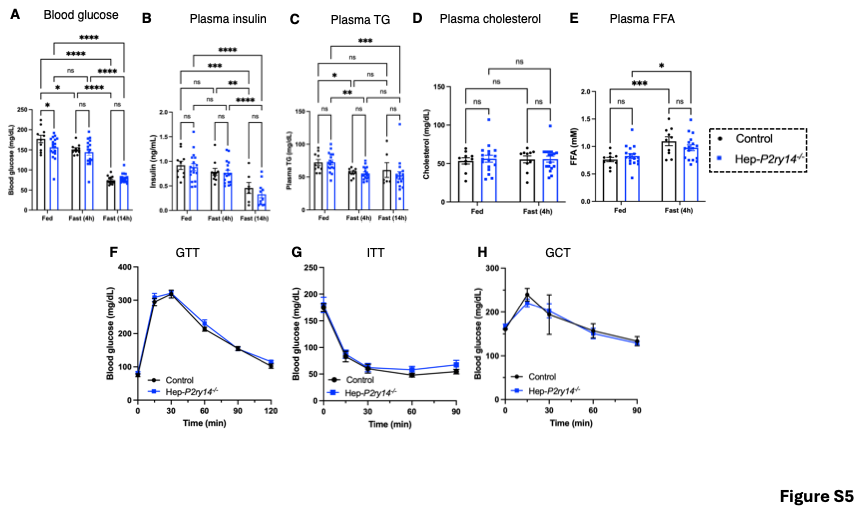

**Figure S5. Hepatocyte-specific deletion of P2Y_14_R does not alter glucose or lipid homeostasis in male mice fed a regular chow diet.** Metabolic parameters were compared between male Hep-*P2ry14*^⁻/⁻^ mice and control littermates. (A) Blood glucose and (B) plasma insulin levels were measured under fed, 4-h fasted, and 14-h fasted conditions. (C) Plasma triglyceride, (D) plasma cholesterol, and (E) plasma free fatty acid (FFA) levels were measured under fed and 4-h fasted conditions. (F) Intraperitoneal glucose tolerance test (ipGTT), (G) insulin tolerance test (ITT), and (H) glucagon challenge test (GCT) were performed in Hep-*P2ry14*^⁻/⁻^ mice and control littermates. Data are presented as mean ± SEM (n = 5–18 mice per group). Statistical significance is indicated as *P < 0.05, **P < 0.001, ***P < 0.0001, and ****P < 0.00001.

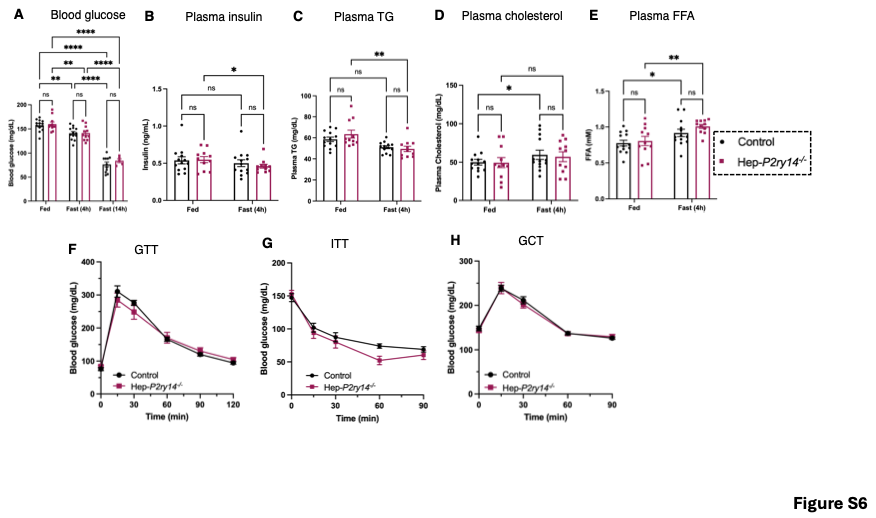

**Figure S6. Figure S4: Hepatocyte-specific P2Y_14_R deletion has no major effect on glucose and lipid homeostasis in female mice maintained on a regular chow diet.** Metabolic phenotyping was performed in female Hep-*P2ry14*^⁻/⁻^ mice and their control littermates. (A) Blood glucose levels were determined in the fed state and after 4 h and 14 h of fasting. Plasma (B) insulin, (C) triglyceride, (D) cholesterol, and (E) free fatty acid (FFA) levels were assessed under fed and 4-h fasted conditions. (F) Intraperitoneal glucose tolerance test (IPGTT), (G) insulin tolerance test (ITT), and (H) glucagon challenge test were performed to evaluate systemic glucose regulation and hormone responsiveness. Data are presented as mean ± SEM (n = 4–14 mice per group). Statistical significance is indicated as *P < 0.05, **P < 0.001, ***P < 0.0001, and ****P < 0.00001.

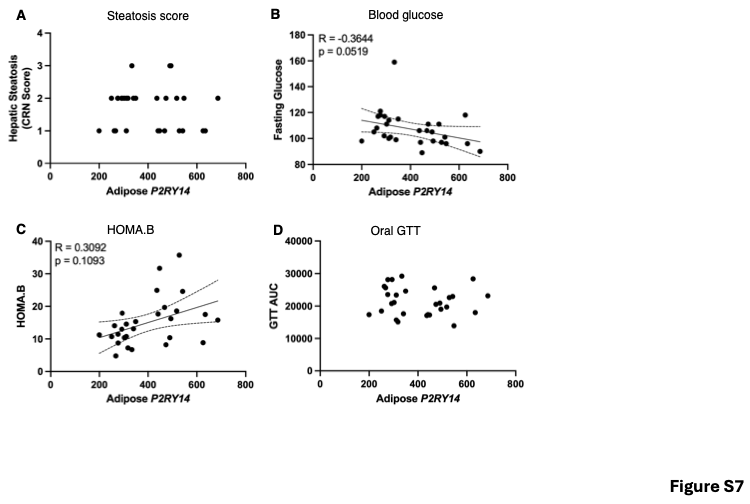

**Figure S7. Correlation between adipose *P2RY14* expression and hepatic steatosis, markers of pancreatic beta-cell function, and insulin resistance.** Adipose tissue samples were obtained from human subjects with MASLD. **(A)** Adipose *P2RY14* expression was not associated with hepatic steatosis, as assessed by the NASH-CRN histological score. Adipose *P2RY14* expression was **(B)** negatively correlated with fasting blood glucose levels, **(C)** positively correlated with HOMA-B, a marker of pancreatic beta-cell function, and **(D)** not correlated with GTT. RNA expression levels were determined by RNA-seq, and correlations were assessed using Pearson’s correlation analysis. MASLD, metabolic dysfunction-associated steatotic liver disease; HOMA-B, homeostasis model assessment of beta-cell function; GTT, glucose tolerance test; NASH-CRN score, NASH Clinical Research Network score.

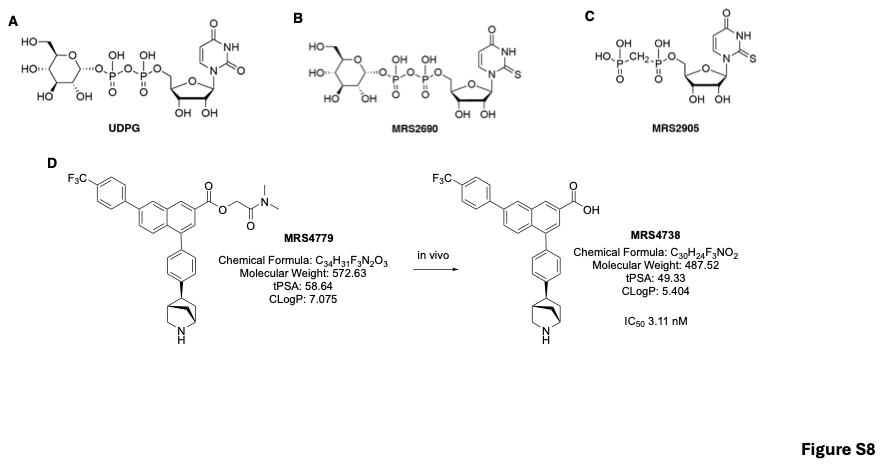

**Figure S8. Chemical structures of P2Y_14_R agonists and an antagonist mono-prodrug.** Chemical structures of **(A)** UDP-glucose, an endogenous P2Y_14_R full agonist (EC_50_=400 nM), **(B)** MRS2690, a UDP-glucose analogue and potent synthetic P2Y_14_R agonist (EC₅₀=49 nM) and, **(C)** MRS2905, a UDP analogue and highly selective and potent synthetic P2Y_14_R agonist (EC₅₀=0.92 nM). **(D)** Chemical structures of the antagonist mono-prodrug MRS4779 and its active drug, MRS4738, a potent P2Y_14_R antagonist (IC₅₀=3.11 nM). MRS4779 is enzymatically cleaved *in vivo* to release MRS4738, thereby improving the physicochemical properties of the active drug.

**Supplementary Table 1. Summary of retention time and transitions for used analytes. Analyte density and quantification of analytes was verified using internal standard (IS).**

| **Compounds** | **Retention time (min)** | **Transition** |
| --- | --- | --- |
| UDP-galactose | 5.80 | 565.09/323.13 |
| UDP-glucose | 6.00 | 565.09/323.13 |
| MRS2905 | 1.00 | 417.04/156.90  417.04/399.00  417.04/271.00 |
| UDP-^13^C_6_-glucose (IS) | 6.00 | 571.21/323.15 |

**Supplementary Table 2. Antibodies used in this study.**

| **Reagent** | **Source** | **Identifier** |
| --- | --- | --- |
| **Antibodies** (Dilution used in parentheses) | | |
| SAPK/JNK Antibody Rabbit mAb | Cell Signaling Technology | 9252 (1:1000) |
| Phospho-SAPK/JNK (Thr183/Tyr185) (81E11) Rabbit mAb | Cell Signaling Technology | 4668 (1:1000) |
| ß-Actin (13E5) Rabbit mAb | Cell Signaling Technology | 4970 (1:3000) |
| ß-Tubulin (9F3) Rabbit Monoclonal Antibody | Abcam | 2128 (1:3000) |

**Supplementary Table 3. Primer sequences (mouse) used for real-time RT-qPCR experiments.**

| **Target Gene** | **Forward primer sequence** | **Reverse primer sequence** |
| --- | --- | --- |
| *P2ry14* | 5’-AGCAGATCATTCCCGTGTTGT | 5’-AGCCACCACTATGTTCTTGAGA |
| *G6pc* | 5’-AGGTCGTGGCTGGAGTCTTGTC | 5’-GTAGCAGGTAGAATCCAAGCGC |
| *Pck1* | 5’-GGCGATGACATTGCCTGGATGA | 5’-TGTCTTCACTGAGGTGCCAGGA |
| *36b4* | 5’-ATG GGT ACA AGC GCG TCC TG | 5’-GCC TTG ACC TTT TCA GTA AG |
